# Adaptation to complex environments reveals pervasive trade-offs and genetic targets with pleiotropic effects

**DOI:** 10.1101/2024.01.24.577006

**Authors:** Alexandre Rêgo, Dragan Stajic, Carla Bautista, Sofia Rouot, Maria de la Paz Celorio-Mancera, Rike Stelkens

**Affiliations:** Department of Ecology and Genetics, Uppsala University, Uppsala 752 36, Sweden; Department of Zoology, Stockholm University, Stockholm 114 19, Sweden; Institute for Fish and Wildlife Health, University of Bern, Bern 3012, Switzerland; Institut de Biologie Intégrative et des Systèmes (IBIS), Université Laval, Québec, QC, Canada; Département de Biologie, Faculté des Sciences et de Génie, Université Laval, Québec, QC, Canada; SciLifeLab, Department of Gene Technology, KTH Royal Institute of Technology, Stockholm 171 65, Sweden; Department of Ecology, Environment, and Plant Science (DEEP), Stockholm University, Stockholm 106 91, Sweden

**Author notes:** Corresponding Author: Alexandre Rêgo. Shared first-authorship (Alexandre Rêgo, Dragan Stajic).

**Keywords:** evolve and resequence, Saccharomyces cerevisiae, complex environments, phenotypic evolution, trade-offs, genomic evolution

## Abstract

Much of our knowledge on the dynamics of adaptation comes from experimental evolution studies that expose populations to a single selective pressure. However, populations in nature rarely adapt to a single stress at a time. Instead, various biotic and abiotic factors come together to produce complex selective environments. Here, we used experimental evolution to describe adaptive dynamics in the presence of simultaneous stressors, and to quantify the evolution of trade-offs between stressors in a complex environment. We adapted populations of the yeast *Saccharomyces cerevisiae* to a full-factorial combination of four stressors over the course of 15 serial transfers. Rapid fitness increases were accompanied by the accumulation of mutations in genes and pathways related to specific stressors. Trade-offs evolved rapidly, with the order and mode of trade-off evolution varying between environments due to the inherent physiological and genetic basis of resistance to each stressor. Compensatory evolution for maladaptation as a result of initial trade-offs was typically mediated by fine-tuning of genes associated with the initially adapted environmental component, rather than those associated with the maladapted trait. As environmental complexity increased, mutations had increasingly broader effects across multiple biological processes. Although shared mutations at the individual SNP level were rare, recurrent mutations affecting the same genes and putative biological processes were abundant across environmental complexity. Our results suggest that adaptation in complex environments follows genetic trajectories shaped by pleiotropy and trade-offs, but these trajectories may impose lasting constraints on future adaptation.

## Introduction

Natural habitats are rich with ecological complexity, resulting in many co-occurring sources of selection through biotic and abiotic factors (Orsini, Spanier, and DE Meester 2012). The dimensionality of concurrent selective pressures may even increase for populations in the Anthropocene, due to additional effects of pollution, urbanization, and climate change (Hendry, Gotanda, and Svensson 2017). Predicting the dynamics and outcomes of adaptation to complex environments has been a long-standing goal with far-reaching implications, from conservation to the evolution of antimicrobial drug resistance (Hendry, Gotanda, and Svensson 2017; Sommer, Johansen, and Molin 2020). But measuring and describing adaptation in the wild is not trivial given logistical limitations to working with natural populations and the difficulty of assigning causes of selection in nature (Vandenkoornhuyse et al. 2010). Experimental evolution studies are now more often implementing temporally and spatially variable or multidimensional selection regimes as a bridge to investigate evolutionary dynamics in ecologically complex natural environments (Barrett, MacLean, and Bell 2005; Kram et al. 2017)(Hendry, Gotanda, and Svensson 2017; Sommer, Johansen, and Molin 2020)(Barrett, MacLean, and Bell 2005; Kram et al. 2017). Even so, few current studies integrate genomic sampling and time-series sampling to address the underlying genomic basis and dynamics of adaptation to complex environments.

Adaptation to complex environments is consistently marked by the existence of trade-offs (Garland, Downs, and Ives 2022). Theory predicts that a particular phenotype cannot be optimal for all selective pressures, and that most populations exist along a range of phenotypes that optimize fitness in multidimensional selection regimes (Shoval et al. 2012). Trade-offs for multiple traits, through resource limitation, antagonistic pleiotropy, genetic correlations, or epistasis are common (Roff and Fairbairn 2007; Garland, Downs, and Ives 2022). However, several experimental evolution studies show that populations can improve fitness in many dimensions simultaneously (Bennett and Lenski 2007; McGee et al. 2016; Bono et al. 2017). Empirical evidence of how trade-offs themselves evolve, and affect dynamics of adaptation over short time scales, is relatively scarce (Roff and Fairbairn 2007; Vincent et al. 2013; Bono et al. 2017).

Adaptation in complex environments is not only influenced by the prevalence of trade-offs. The dynamics of phenotypic evolution (i.e. the rate of fitness changes and trait trajectory) and genomic evolution (i.e. the rate of allele frequency change, and the number of adaptive mutations) can also be affected by the complexity of the environment. For example, when multiple stressors simultaneously increase the strength of selection, the trajectory and rate of trait evolution can be altered (H. A. Orr 2005; Cisneros-Mayoral et al. 2022). Increased selection in more complex environments may also have demographic effects. Reduced population sizes can lead to reduced genetic diversity and mutational supply, as well as increased genetic drift (Ewens 1979). Furthermore, adaptation to multiple concurrent stressors may require distinct solutions that differ from those needed for individual stressors (Elena and Lenski 2003).

Many studies focus on adaptive outcomes and trade-offs, but only a few use time-series data to describe the dynamics of trade-off evolution itself (Huang et al. 2017; J. A. Orr et al. 2022). To understand how the dynamics and outcomes of adaptation differ between simple and complex environments, we used experimental evolution with the yeast *Saccharomyces cerevisiae.* Here we define simple environments as those manipulated along a single experimental dimension and complex environments as those manipulated along multiple dimensions, while acknowledging that even “simple” environments contain numerous factors that cannot be completely isolated or controlled. We evolved replicate lines in four single stressors and full-factorial combinations of the four stressors. We collected fitness and genomic time-series data to investigate the tempo and mode of adaptation at different biological scales, including trajectories in multidimensional adaptive space that populations explore when environmental stressors coincide. To test for the existence and genetic basis of trade-offs, we measured fitness and sequenced genomes of populations that had evolved in complex environments. We tested these populations in both the original complex selective media and in simpler media containing only one component at a time. Our goals were to determine (i) how the dynamics of adaptation depend on the complexity of the environment, (ii) how does the genetic basis of adaptation differ in complex environments (iii) to what extent trade-offs exist in complex environments, and (iv) how do these trade-offs evolve.

## Results

Initially isogenic replicate lines of *S. cerevisiae* (four replicates per environment, 60 in total) were evolved in either single stressors (Figure 1A, 1D environments, designated with single-letter abbreviations) or the full-factorial combinations of multiple stressors (Figure 1A, 2D-4D environments, designated with multi-letter abbreviations). The four stressors used were high salinity (0.8M NaCl, designated “N”), the antifungal drug fluconazole (20μg/ml, “A”), starvation (0.5% glucose YPD, “S”), and high temperature (37°C, “H”). The selective pressure of each environment was quantified at the beginning of the experiment as the difference in final optical density (OD600) after 24 hours relative to growth in a benign YPD environment (ΔOD=OD_YPD_ - OD_Selected_, Mean±SE) (Figure 1B, see Figure S1 for stressor effects based on growth rate). Of the 1D environments, the salt (ΔOD = - 0.520±0.006) and starvation environments (ΔOD = -0.399±0.008) had the largest stressor effects, while antifungal (ΔOD = -0.143±0.159) and heat (ΔOD = -0.116±0.048) had more moderate effects (Figure 1B). Experimental evolution on each line was conducted for a total of 15 transfers with transfers occurring every 48 hours into fresh media (Figure 1C). Evolving lines were frozen at the midpoint of the experiment (T7) and the end (T15) for further genomic sequencing.

**Figure 1.**
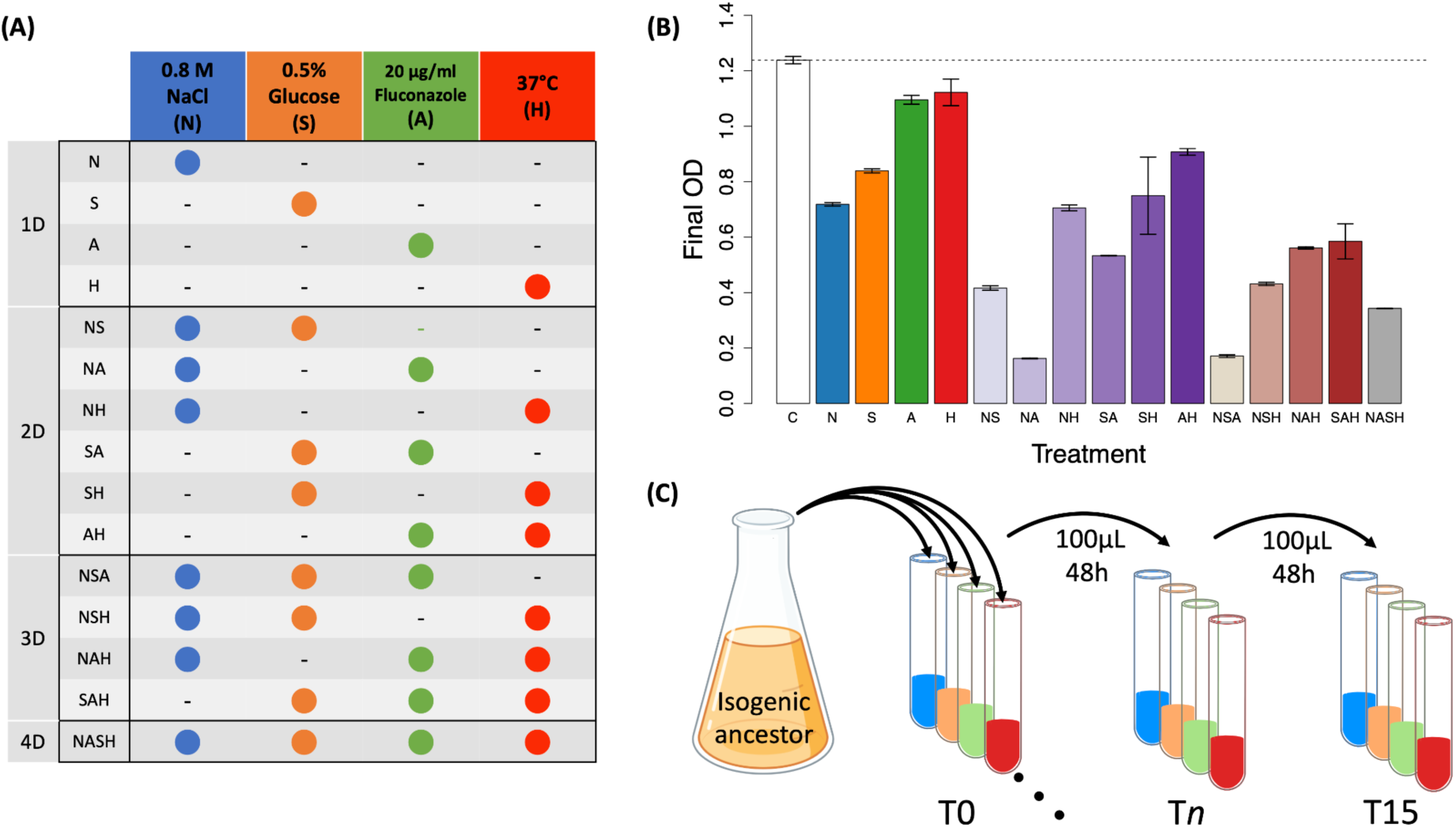
Experimental set-up and stressor effects for each environment. (A) Populations were evolved in every stressor and full-factorial combination of each. Colored dots represent the presence of a stressor. Abbreviations are given for each combination of stressors, as well as the ‘dimensionality’ (1D-4D) of the environmental complexity. (B) The stressor effect of each environment in the ancestor was measured as the difference in final OD after 24 hours in a selective environment relative to benign YPD medium. The mean estimate is given with standard error bars. (C) From an initially isogenic colony, four replicate populations of yeast were inoculated into each environment. Populations were then evolved through serial transfer every 48 hours where 2% (100μL) of the culture was transferred into fresh media, for a total of 15 transfers. Fitness measurements and whole genome, whole population sequencing were conducted at T7 and T15.

### Rapid but plateauing adaptation to environmental stressors

To determine how much a replicate line had adapted, we measured the fitness of evolved lines relative to the fitness of the ancestral population in growth competition assays, designated as the selection coefficient (Gordo, Perfeito, and Sousa 2011). All lines showed evidence of adaptation at the midpoint of experimental evolution (T7), with few exceptions (all four replicates of SH, and NASH1) (Figure 2). The largest selection coefficient at the midpoint was observed in the antibiotic environment (*S*_*A*2_=0.43). However, SA lines showed the highest selection coefficients on average (<u>*S*</u>_SA_=0.41, SE=0.007). By the midpoint of the experiment, the fitness of many populations had reached a fitness plateau after which fitness did not significantly increase again until the end of the experiment (paired *t, p*>0.05), with some exceptions (average of replicates for the N, NA, NH, NAH, SAH, and NSA lines). Despite many populations not differing significantly in fitness between T7 and T15, average gains in fitness between T7 and T15 did tend to increase as complexity levels increased (β = 0.024, t(58) = 2.06, p = 0.044), indicating that populations in more complex environments had not yet reached their local fitness optima.

**Figure 2.**
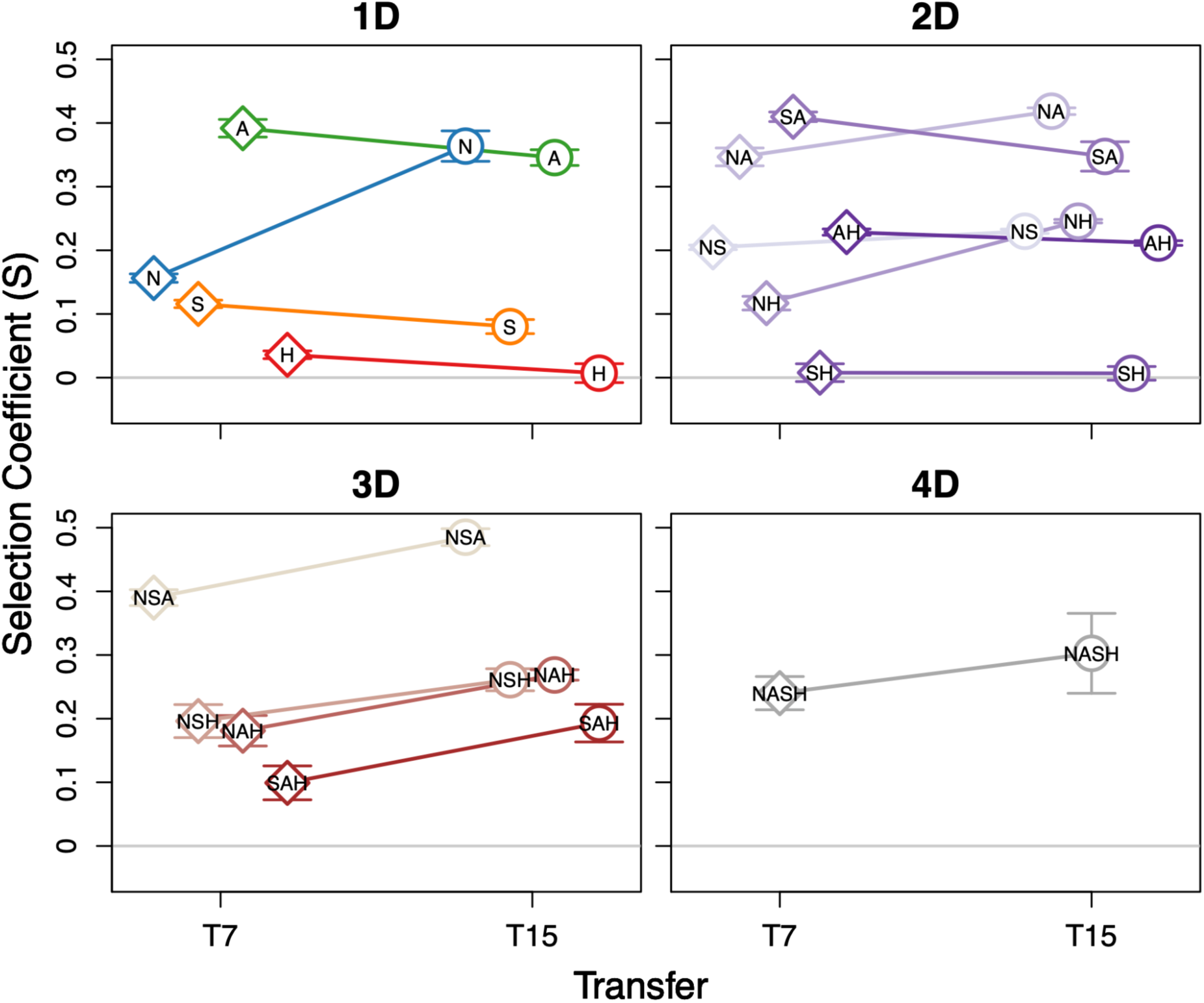
Selection coefficients at T7 and T15 for each environment, partitioned by environmental complexity level (1D–4D). Mean selection coefficients of the four replicates for each treatment are given as points while bars represent the standard deviations. T0 is excluded for visualization purposes, but would reside on the S=0 line.

### Adaptation to complex environments reveals pervasive trade-offs

We tested for the existence of trade-offs during adaptation to the individual stressors (N, A, S, H) of the complex environments. For this, we compared the fitness of evolved lines adapted to complex environments to the fitness of single-stressor specialists in the respective single stress environment. Populations evolved in complex environments were unable to adapt to the same level of fitness as all single environment specialists in their respective environments, resulting in abundant trade-offs (Figure 3, Figure S2). For each of the four individual stressors, there were 7 complex environment combinations which contained the stressor in our full-factorial design. The individual stressor with the highest prevalence of trade-offs (i.e. the proportion of complex environment-adapted lines which on average were unable to reach the fitness of single stressor specialists; *t*, *p*<0.05) was NaCl (6/7 complex-adapted lines), followed by Starvation (3/7), Heat (3/7), and Antifungal (1/7) (Figure S2, Figure S3).

**Figure 3.**
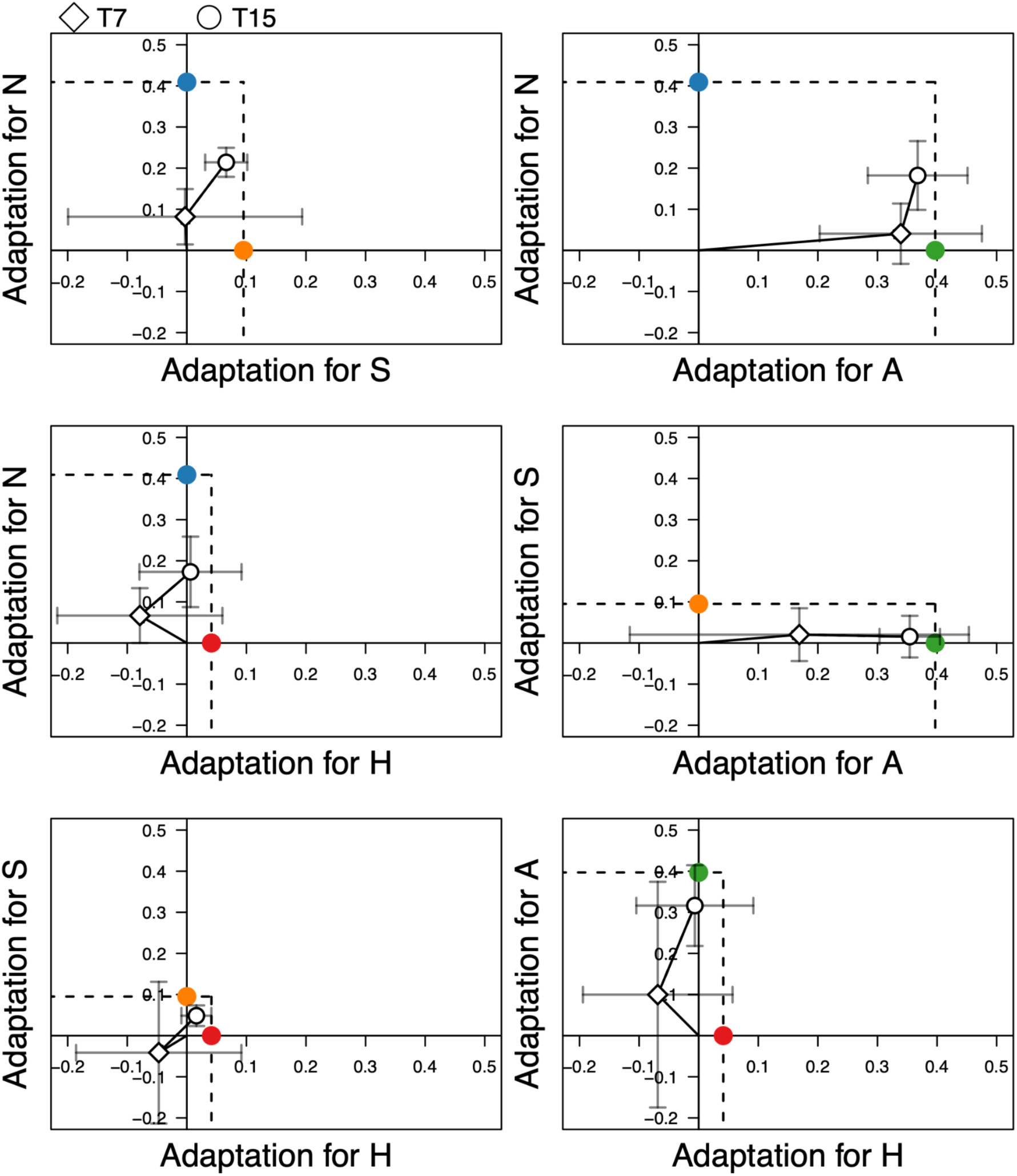
Trade-offs in adaptation to individual stressors within complex environments. Each panel shows adaptation (relative fitness compared to ancestor) of populations evolved in complex environments when tested in single-stressor conditions. For each pairwise comparison, means (and ±1 sd bars) are shown for those complex environments that contain both individual stressors as diamonds (at T7) and circles (at T15). Colored dots represent the fitness of single-stressor specialists projected onto their respective axes. The lower-left area of the black dotted lines represents the space in which populations evolved in complex environments are expected to fall (i.e. less adapted in single-stress environments than either 1D specialist).

To evaluate how well multi-stressor adapted lines (e.g., NAS lines) performed compared to single-stressor specialists (e.g., N, A, and S lines), we calculated their relative Fitness 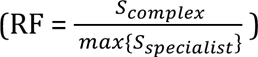 in each respective environmental component. The evolution of trade-offs under heat stress showed a particularly strong initial fitness cost at T7 (<u>*RF*</u> = - 1.60) which was later mitigated by T15 (<u>*RF*</u> = 0.05). Lines tested in other single stressors experienced less severe trade-offs, with <u>*RF*</u> gradually increasing: from 0.54 to 0.87 for antifungal, from 0.21 to 0.47 for salt, and from -0.04 to 0.43 for starvation (Figure S3). These results suggest that, when present, populations typically adapt to antifungal first, reaching a greater relative fitness than when adapting to any other stressors by T15. The adaptation of populations to antifungal seemed to incur negative trade-offs to heat stress, though these maladaptations were typically compensated for by T15 (Figure S3). Surprisingly though, some lines evolved in complex environments were able to exceed the fitness of the single-stressor specialists. This is especially true for NAH lines tested in antifungal medium and NS lines tested in starvation medium, where all replicates showed higher fitness than the specialists in the specialist’s respective stressors (Figure S3, p<0.05).

### Trade-off and cross-protective evolution to non-selected environments

To test for possible novel trade-off or cross-protective mechanisms between adaptation to different stresses, we tested the fitness of each population (all 1D–3D populations) in every respective environment not directly selected for during experimental evolution (e.g. N-evolved populations would be tested in A, S, and H environments individually, Figure 4). The strongest trade-offs across all evolved populations were observed in novel antifungal exposure, where the majority of evolved lines (6/7) showed a significantly reduced fitness in the presence of fluconazole at T7 (Figure 4, upper right panel; <u>*S*</u>_*A_T7*_= -0.24, SD = 0.19). While this sensitivity was reduced by the end of the experiment, it rarely led to a fitness advantage ( <u>*S*</u>_*A_T*15_ = -0.16, SD = 0.16). In contrast, we often observed significant cross-protective responses in other non-selected environments by T15 (2/7 in S, 4/7 in N, and 5/7 in H) . This suggests the existence of cross-talk between stress response pathways to these different environmental stressors. We observed the largest cross-protection in the case of exposure to osmotic stress at T15 (<u>*S*</u>_*N_T*15_=0.049, SD = 0.09; Figure 4, upper left panel), where surprisingly even populations adapted to antifungal showed an increase in fitness, despite low fitness of N-evolved populations tested in the antifungal environment.

**Figure 4.**
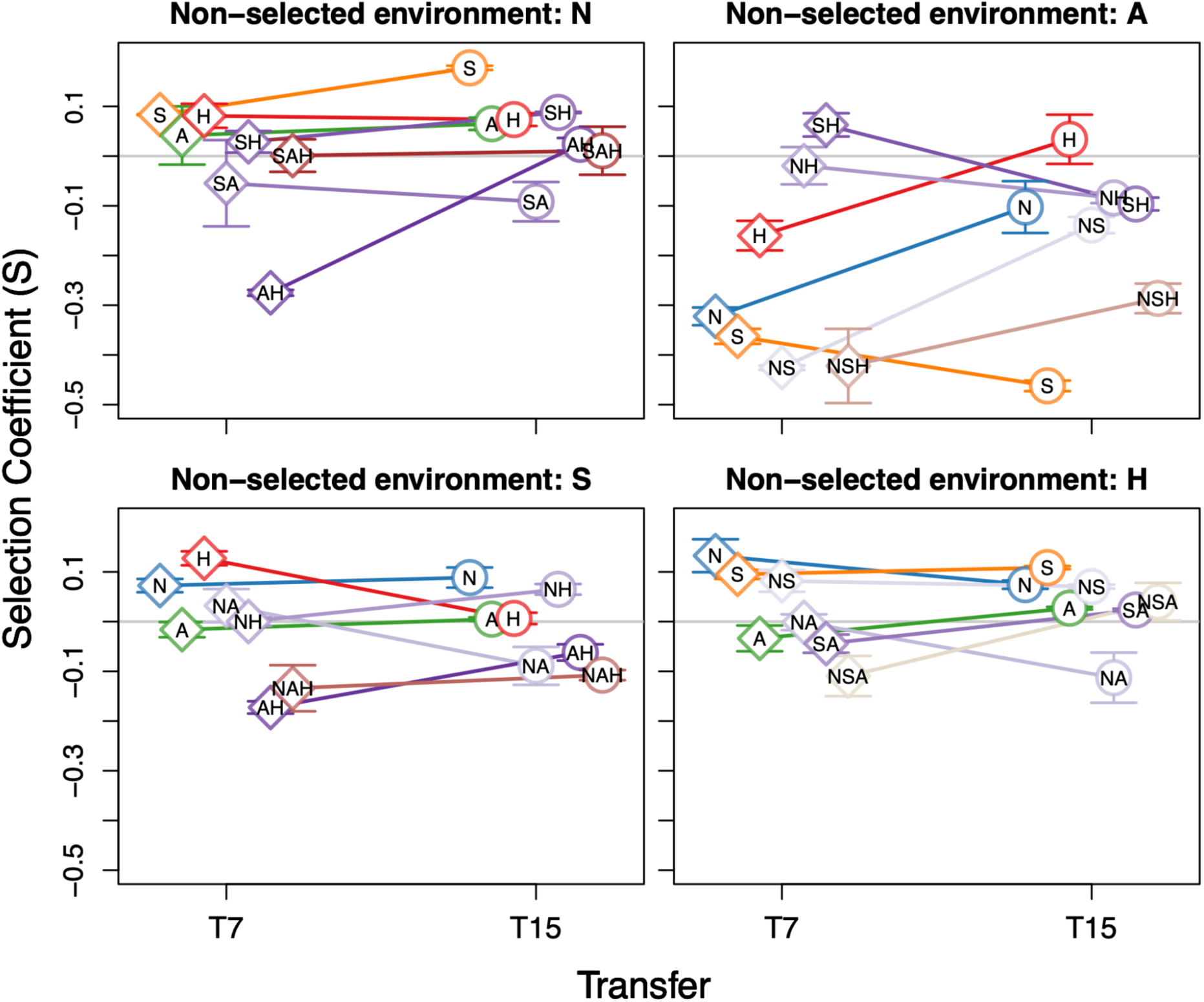
Patterns of trade-offs and cross-protection evolution in non-selected environments. Estimated selection coefficients are shown as diamonds or circles for T7 and T15, respectively, with bars as standard deviations. The grey line represents a selection coefficient of 0.

We tested whether the degree of environmental complexity a line evolved in (1D-3D) had a bearing on the degree of cross-protection to each novel environment (except NASH replicates as they were exposed to all stressors) with linear regression. By the end of experimental evolution, we found a negative association between fitness and complexity in the presence of salt and starvation (*p* < 0.05), whereas no association was found in either antifungal or heat (Figure S4).

### Fitness costs of adaptation

We tested for possible costs of adaptation to stressful environments by measuring the selection coefficient in the originally benign YPD environment for each evolved population at both time points (*S_YPD_Tn_*). Across all treatments and lines, we observed an average decrease in fitness in YPD at T7 (<u>*S*</u>_*YPD_T7*_ = -0.022, SD= 0.13). However, this cost diminished (or was reversed) by T15 of the experimental evolution (<u>*S*</u>_*YPD_T*15_ = 0.013, SD = 0.066; paired *t* = -2.62, p = 0.011) (Figure 5), indicating the evolution of cost compensatory mechanisms. We observed higher mean adaptation costs in more complex environments (>1D) relative to single stressor environments (1D) at both T7 (*t* = 5.23, df = 54.544, *p*<0.01) and T15 (*t* = 4.26, df = 43.714, *p* < 0.01). The fitness costs associated with adaptation varied among stressors and across time points. At T7, adaptation to heat stress imposed the highest fitness cost in YPD (F(1,55) = 7.93, p < 0.01), while at T15, adaptation to antifungal resulted in the greatest fitness cost in YPD (F(1,55) = 4.77, p = 0.033).

**Figure 5.**
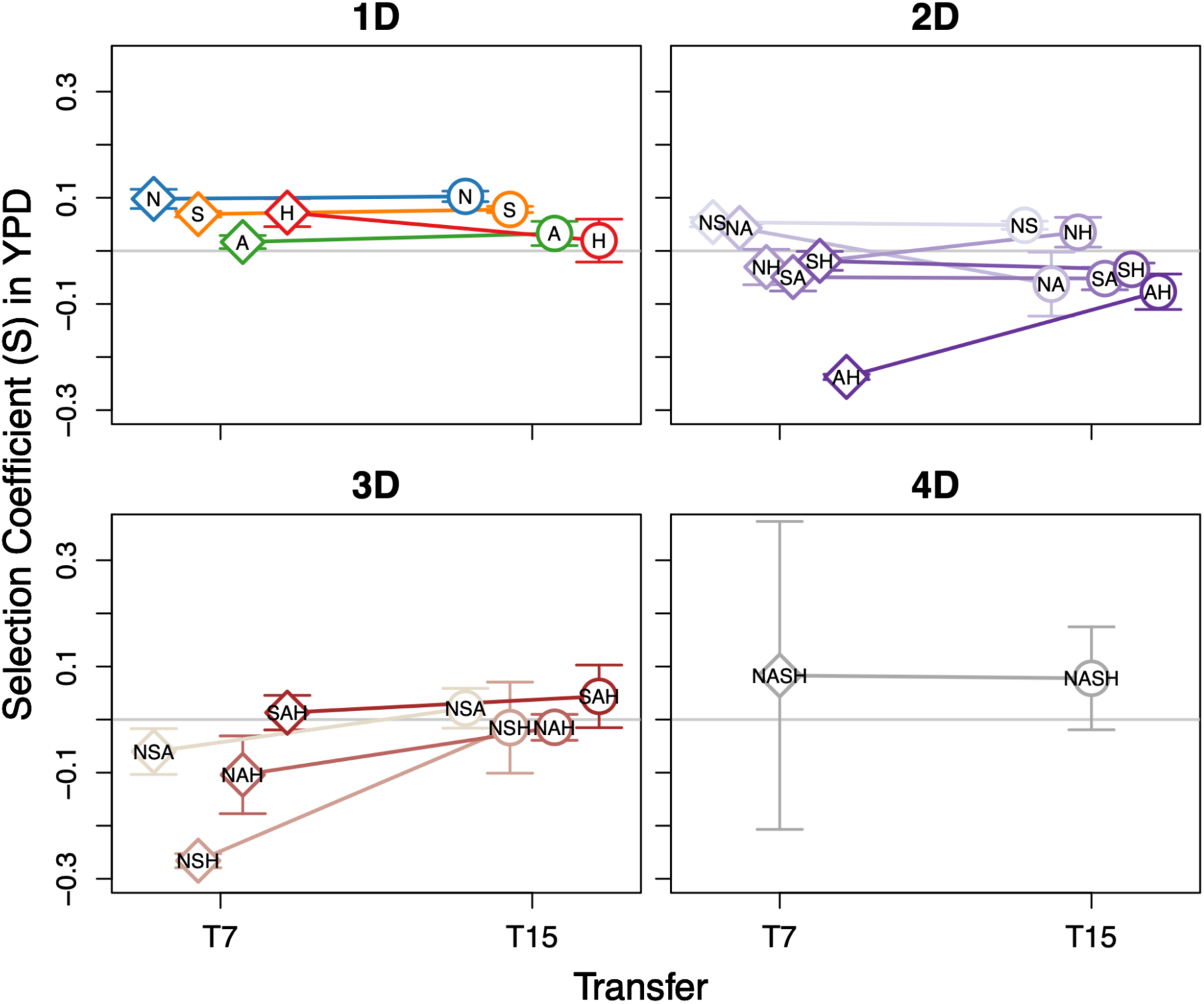
Cost of adaptation in the ancestral environment (YPD) for each evolved population, partitioned by environmental complexity level (1D–4D). Estimated selection coefficients are shown as diamonds (T7) and circles (T15) with bars as standard deviations. The grey line represents a selection coefficient of 0. T0 is excluded for visualization purposes.

### Genomic dynamics and basis of adaptation characterized by few, but pleiotropic, mutations

We identified 228 SNPs and 57 INDELs across all timepoints and populations in our experiment. Populations varied moderately in SNP and INDEL number by T15 (mean = 3.91, SD=3.55), with some populations seemingly containing no mutations (see Discussion) while the highest number of mutations (n=18) was in the antifungal-adapted population, A2. Despite clear instances of phenotypic evolution in most populations (Figure 2), mutational sweeps were rare. By T7, 21.6% (13/60) of populations had at least one mutation fixed or nearly fixed with an allele frequency greater than 0.80, and this proportion increased to 31.6% (19/60) by T15 (Figure 6). The probability of observing sweeps by T15 was stressor-biased, where sweeps were more associated with evolution in antifungal (*β*=0.676, p<0.001) than salt and starvation (both *β*=0.363, p=0.037), and heat (*β*=0.176, p=0.305). Although we do not formally quantify selection (due to hitchhiking mutations) at the genomic level, predicted effects of mutations by snpEff showed that the mutations that did sweep were primarily mutations of at least moderate putative effect (Figure S5). We additionally observed some signals of clonal interference, especially between the mutations conferring antifungal resistance, where initially high frequency mutations (at T7) had decreased in frequency by T15, likely in response to the acquisition of a new mutation (e.g. A3, NA3, NASH4). Of the 129 high frequency mutations at the midpoint of the experiment, 83 (64%) mutations across 32 (of 60) lines decreased in frequency by at least 10% by the end of the experiment.

**Figure 6.**
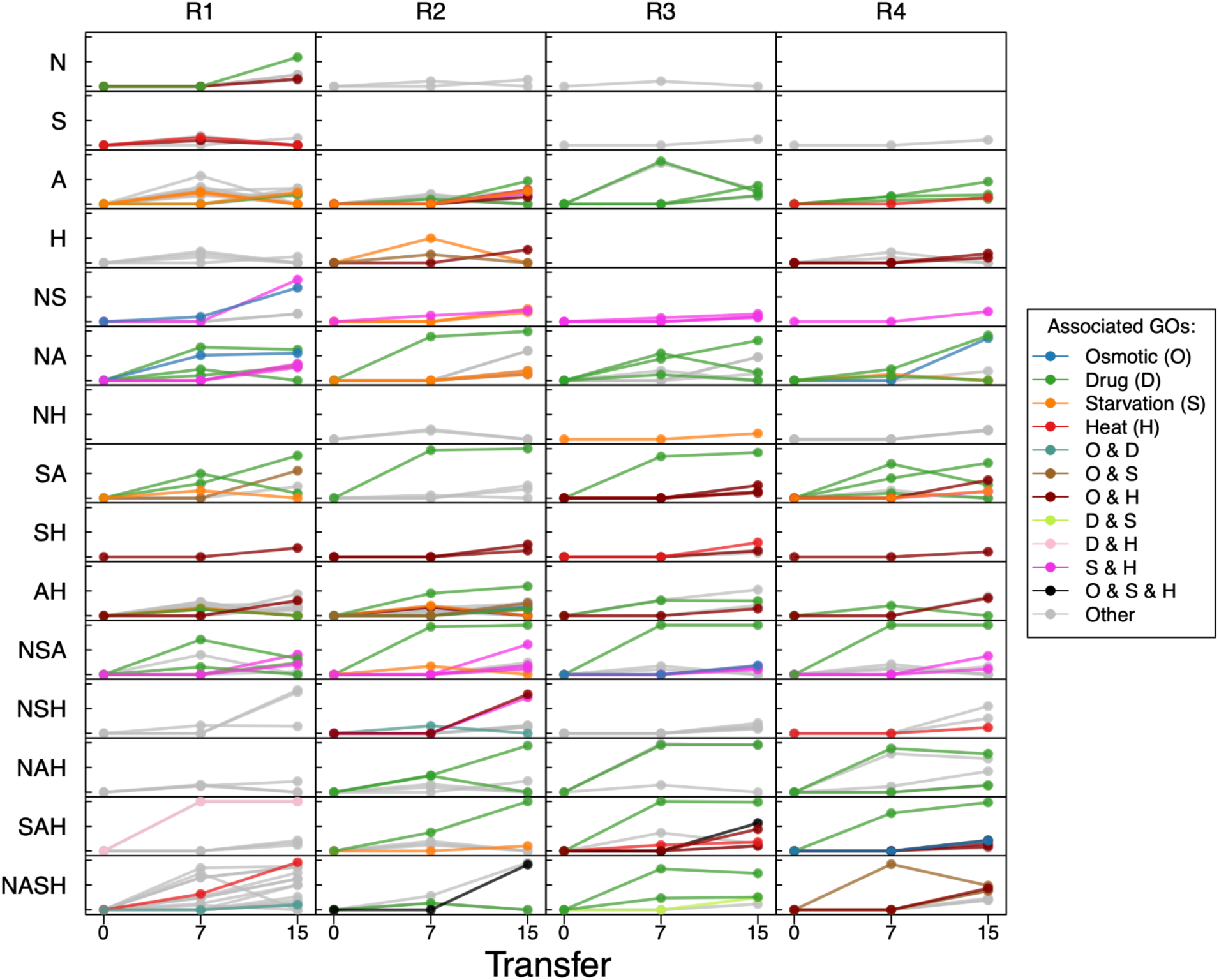
Allele frequency dynamics in evolved lines. Points represent allele frequencies at time points at T0, T7, and T15. Points are colored by association with general gene-ontology terms: Response to osmotic stress (O), response to starvation (S), drug response (D), and response to heat (H). Some mutations have overlapping GO terms, and are colored separately for each type of overlap. Replicates are designated as R1–R4.

We also found larger structural variants during experimental evolution by identifying chromosomes or regions within a chromosome with elevated coverage relative to the genome-wide or chromosome-level sequencing depth, respectively. At the chromosome level, we observed that chromosome 12 showed consistent patterns of copy number gain (aneuploidy) across several line replicates (NSA1, NSA3, NAH1, SAH1, NASH1) while chromosome 14 was elevated in one replicate (also in NAH1) (Figure S6). Chromosome 12 contains genes related to pleiotropic drug resistance (PDR8, TOP1, CIS1), osmotic stress (FPS1, RCK2, STE11, SGD1, SSK1), glucose limitation (SSA2, YPT2, GSY2, ECM30, FBP1), and heat stress (SSQ1, SYM1, GSY2, HSP60,HSP104, SSA2). Chromosome 14 contains genes for osmotic stress (VAC7, POR1, SSK2, INP52,SKO1, FIG4), pleiotropic drug response (MKT1,HCH1, PDR16,HDA1), and heat stress (HCH1, WSC2, HDA1, YDJ1, NMA111).

Within individual chromosomes, only two partial CNVs were identified (Figure S7). The first in the NAH1 population between 90kb–150kb on Chromosome 3 (spanning 35 genes) at both T7 and T15, and the second in SAH2 only at T7, between 46kb–70kb on chromosome 15 (spanning 14 genes). The partial CNV in NAH1 contains genes related to salt homeostasis (SAT4, RVS161), and the CNV in SAH2 contains genes related to glycolysis (PFK27) and transcriptional response to glucose starvation (RTC1), among others.

We found that populations typically contained mutations whose putative effects (based on gene ontology terms, see Methods) matched their evolved environment (Figure 6). In particular, we found that almost all populations evolving in the presence of antifungal (27/32) acquired mutations in at least one of three genes (PDR1, PDR3, and PDR5) encoding for pleiotropic drug response factors or transporters (Akache et al. 2004) and are known to confer resistance to fluconazole (Spampinato and Leonardi 2013). Furthermore, antifungal-related mutations are often the first to appear and rise in frequency in these populations, suggesting that adaptation to antifungal is more mutationally accessible (i.e., simpler genetic basis and possibly higher genome-wide mutation rate) than other stressors.

Despite abundant mutations with clear associations to the environment they appeared in, many genes targeted by mutation are likely pleiotropic, affecting many biological processes (Figure 7A). Here we use the number of gene-ontology terms (pruned for redundancy using R package `rrvgo` v1.12.2) for each mutation as a proxy of pleiotropic effects. Strikingly, we observed that the pleiotropy of mutated genes increased as complexity increases (ꞵ=1.162, p=0.028, Figure 7B).

**Figure 7.**
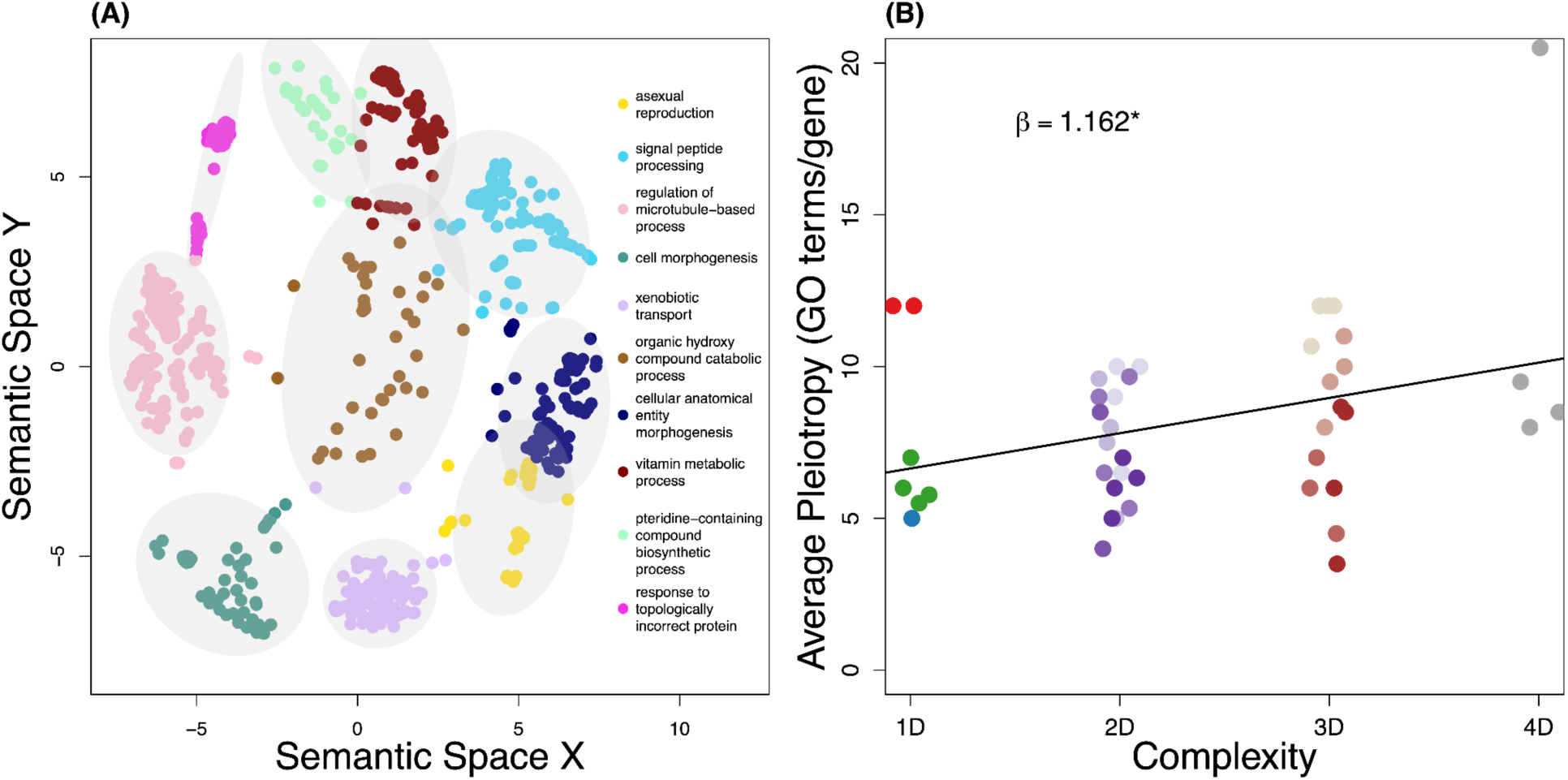
Pleiotropy of genes mutated during experimental evolution. (A) Visualization of the types and number of biological processes associated with all mutations observed during experimental evolution. Gene ontologies were pruned for redundant terms based on semantic similarity using the R package rrvgo and clustered for visualization with k-means clustering (k=10). For each cluster, the term with the lowest dispensability score is shown. (B) Average number of GO terms associated with gene mutations in each level of environmental complexity at the end of experimental evolution (T15). GO terms were first clustered by semantic similarity using R package rrvgo to reduce redundancy. To account for mutations which may not contribute to adaptation, we consider only those which have increased to an allele frequency of at least 20% in any population. The slope of the regression is plotted and shown. The asterisk indicates a significant slope.

### Parallel mutational targets across levels of environmental complexity

We observed high genetic parallelism (Figure 8), with 44/60 lines sharing at least one genic mutation with another independent line. These recurrent mutational targets consisted of 20 genes (13.89% of the 144 total genes). The PDR family of genes, involved in pleiotropic drug response, showed the most prominent mutations, with nearly all lines evolved in the presence of fluconazole (27/32) acquiring mutations in either PDR1, PDR3, or PDR5. PUF3 had the second-highest number of shared mutations, primarily appearing in populations exposed to high heat (particularly in all four SH replicates). Notably, PUF3 null mutants are associated with innate heat tolerance (Harris et al. 2005; Jarolim et al. 2013). Although high heat imposed the lowest stressor effect (Figure 1C), heat tolerance may be easily accessible through loss-of-function mutations of PUF3.

**Figure 8.**
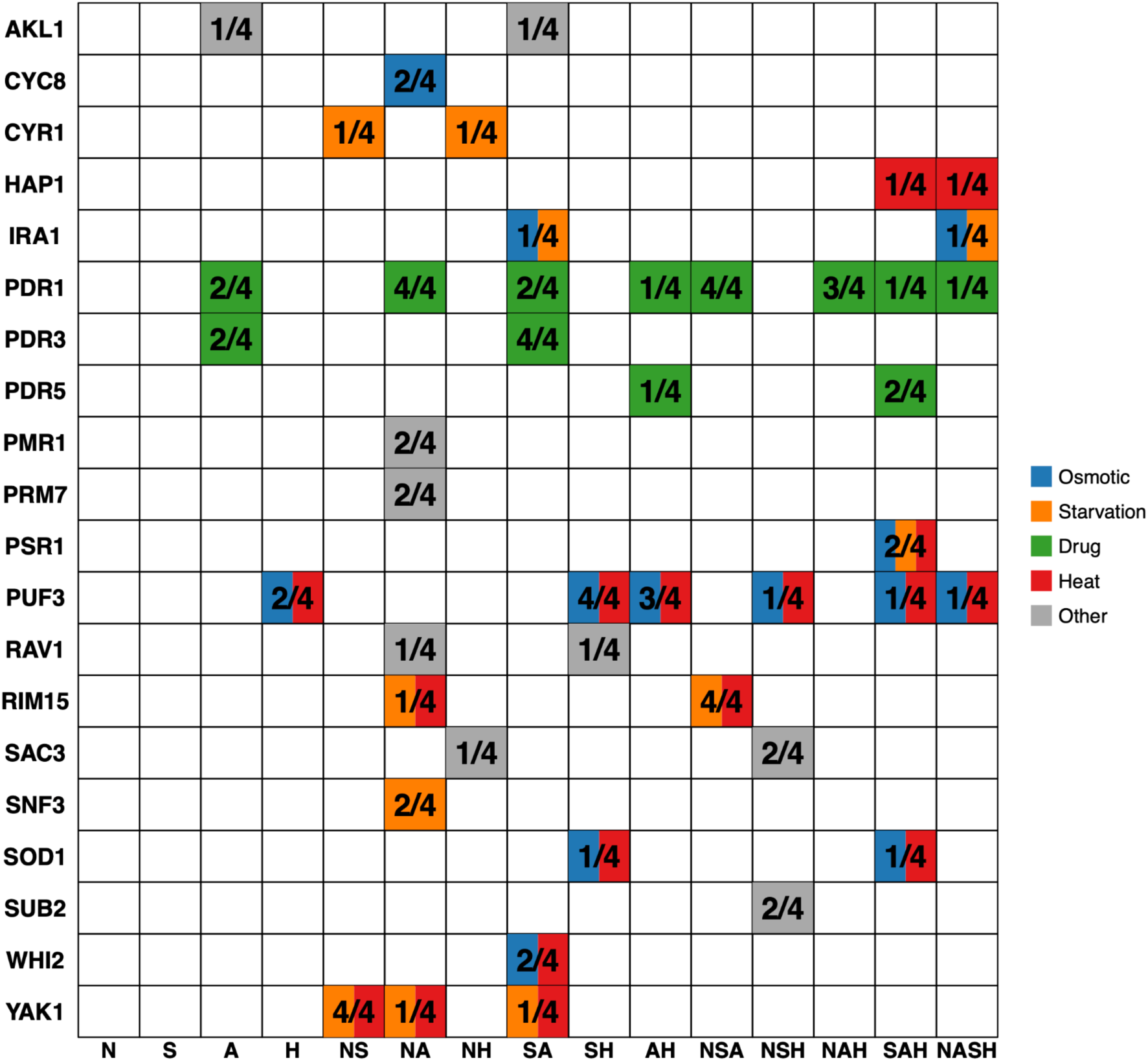
Recurrent mutations in the same gene across independent evolved populations. Shared genic mutations are colored by the parental GO category, white indicates no shared mutation. The fraction indicates the number of line replicates (of four total) which harbor that gene mutation.

At the nucleotide level, 10 point mutations were shared between two or three independent lines by the end of the experiment (Figure S8). Consistent with the genic-level findings, the most common recurrent SNP mutations were in PDR1 and PUF3. Both PDR1 and PUF3 showed recurrent mutations at two different positions. Shared SAC3 mutations, associated with yeast bud and cell size morphological changes (Watanabe et al. 2009; Jorgensen et al. 2002) and possible heat resistance (Jani et al. 2009), were observed in NSH1 and NSH3. We also observed shared mutations in several other genes with known functions: SAC3, associated with bud and cell size morphological changes and increased heat resistance in S288c, appeared in NSH1 and NSH3; YAK1, encoding a regulator of glucose-deprivation stress response (Moriya et al. 2001), in NS1 and NS3; and SOD1, where mutations double yeast lifespan during stationary phase (Harris et al. 2005), in SH2 and SAH4. The remaining shared point mutations occurred in genes of unknown function: MLF3 (in S3 and S4), DFP1 (in NSH2 and NASH4), and YDL228C (in NS1 and SH3).

### Gains and losses of putative compensatory mutations during trade-off evolution

We found several fitness trade-offs at T7 where populations were maladapted to one of the individual components of their complex environment. However, most trade-offs resulting in negative fitness at T7 had been compensated for by T15 such that relative fitness had at least increased to nearly neutral (≈0) or positive values (Figure S2, Figure S3). We further investigated putative mutations concomitant with the largest fitness compensations (Supplementary Table 1).

In lines with the greatest compensatory evolution, fitness changes between T7 and T15 typically were more often concomitant with gaining additional *de novo* mutations rather than the loss of *de novo* mutations. Many of these new mutations were not clearly related to a biological function associated with the tested single-stressor. Additionally, when mutational losses occur, the functions of the affected gene are more likely to be associated with a different stressor than the one tested. For example, the NASH1 line had some of the greatest compensatory evolution in both salt (fitness change from S_T7_=-0.21 to S_T15_=0.12) and high heat (S_T7_=-0.38, S_T15_=-0.14) conditions. While the mutations gained between the midpoint (T7) and end (T15) of experimental evolution were not directly related to either salt or heat stress (affecting protein phosphorylation [SKY1], vacuolar protein catabolism [PRC1], and mitochondrial nuclease and mRNA processing [BI2 and BI3]), the single mutation lost was in CSG2. CSG2 encodes a regulatory subunit for mannosylinositol phosphorylceramide synthase, which works with ergosterol to maintain cell wall integrity. Notably, the mechanism of fluconazole action involves inhibiting ergosterol synthesis, thereby disrupting cell wall permeability (Spampinato and Leonardi 2013). The loss of the CSG2 mutation corresponded with decreased antifungal fitness in NASH1 from S_T7_=0.25 to S_T15_=0.15. Similar patterns were found in lines such as NSA3 (loss of FCY2, which shows cross-resistance to fluconazole) and NSA4 (loss of TEC1, which prevents penetration of fluconazole into the cell wall of *Candida albicans*).

This general pattern of compensatory evolution also occurred in cases not involving antifungal resistance, though to a lesser extent. For instance, the NSH2 line demonstrated some of the greatest compensatory evolution under both glucose starvation (S_T7_=-0.29 to S_T15_=0.19) and heat stress (S_T7_=-0.23 to S_T15_=0.1). NSH2 gained mutations in genes associated with chromosome segregation (OKP1) and mitotic recombination (THP2). While NSH2 acquired mutations potentially relevant to the tested stressors, including PUF3, which has been linked to heat shock response, and IRA2, which regulates cAMP levels during nutrient limitation, NSH2 lost a mutation in AUR1, a gene associated with hyperosmotic stress resistance. Unlike in antifungal populations, the loss of AUR1 did not decrease NSH2’s salt tolerance (S_T7_=0.1 to S_T15_=0.25). These observations support a two-stage evolutionary process in which adaptation occurs initially to a primary stressor that compromises resistance to secondary stressors, followed by evolutionary refinement that enhances overall stress resilience while sometimes reducing the effectiveness of the initial adaptation (as seen with antifungal resistance).

## Discussion

Natural environments in which evolution occurs are immensely complex, with variation across multiple axes and timescales. While experiments in simplified environments in the laboratory have produced key insights about fundamental evolutionary processes and mechanisms, they have been criticized for lacking ecological realism, potentially limiting our understanding of evolution in natural populations (Waldvogel et al. 2020; Bergelson et al. 2021; Venkataram and Kryazhimskiy 2023; Zelnik et al. 2024). As a stepping-stone between simplified laboratory environments and nature, we adapted populations of yeast to a full-factorial combination of four stressors to assess the mode and dynamics of adaptation as environmental complexity increases. Our results demonstrate that adaptation to complex environments imposes trade-offs between traits whose dynamics can rapidly evolve. The order of adaptation to stressors was biased first by mutational accessibility and subsequently by the strength of selection. Trade-offs imposed fitness costs not only in stressors that had been simultaneously selected for, but also in stressors that the evolving lines had never been exposed to, and even in the initially benign ancestral environment. However, initial trade-offs were often compensated through mutational fine-tuning, with mutations of greater pleiotropic effect arising in more complex environments.

### Dynamics and genetic basis of adaptation to complex environments

We observed rapid fitness increases over the course of 15 serial transfers in almost all treatments and lines (Figure 2). While many lines had plateaued by the experiment’s midpoint, those in higher-dimensionality environments continued to gain fitness between T7 and T15, albeit at a reduced rate. This pattern of initially rapid adaptation followed by a fitness plateau aligns with previous studies of microbial evolution from *de novo* mutations (Wiser, Ribeck, and Lenski 2013; Johnson et al. 2021). The continued evolution of improved fitness in complex environments likely stems from two factors. First, populations experiencing multiple stressors may start further away from their fitness optimum as a result of increased environmental stress and require more time and mutational steps to approach the optimum (Hoffmann and Hercus, 2000). Adaptation to complex environments may also have a more polygenic genetic basis (Anderson et al. 2014), requiring additional time to acquire the necessary mutations.

At the genomic level, we observed the appearance and subsequent rise in frequency of SNPs, INDELs, and copy number variants during evolution (Figure 6, Figure S6, Figure S7). Genomic evolution in our lines was characterized by few mutational sweeps and moderate repeatability. The paucity of sweeps may be due to the effects of clonal interference, which frequently occurs in large populations such as in our experiment (Park and Krug 2007), which can slow down the rate of genomic adaptation (Miralles et al. 1999; Hughes et al. 2012). Lines adapting to the presence of fluconazole were notable exceptions, often exhibiting rapid sweeps and parallel gene mutations. This exception may be attributed to the simple genetic basis of antifungal resistance in conjunction with a heightened mutational supply. Not only is fluconazole known to increase the rate of DNA damage (Silva et al. 2019; Yüzbaşioğlu et al. 2008; Ribeiro et al. 2022), but the most common mode of evolution for antimicrobial resistance typically comes from either loss- or gain-of-function mutations in a single gene (Bavro, Conn, and Davies 2019). This is reflected in the recurrent mutations found in the PDR family of genes, in which mutations have been shown to increase the efflux of azoles in other studies of yeast by up to 40 times (Wang et al. 2022).

Many genes targeted by mutations during experimental evolution were associated with multiple biological processes not immediately related to the environmental components. We observed that mutations associated with a greater number of different biological processes arose in environments of higher complexity. This suggests that acquiring mutations with pleiotropic effects, which may perform multiple functions (though less than optimally), results in higher fitness than acquiring multiple mutations in different genes that each affect only one particular trait, or that mechanistic constraints exist between genes of specialized functions. However, since greater pleiotropy was observed only at the end of the experiment, in conjunction with an order bias of adaptation and patterns of compensatory evolution to individual stressors (see below for further discussion), this suggests that adaptation first proceeds by mutations in genes of specialized function, whose functions are fine-tuned by mutations of pleiotropic effect.

### Trade-off evolution between environmental stressors

Fitness trade-offs were abundant across all environments and conditions. Our fitness assays showed that no line evolving in a complex environment was able to reach the same level of fitness in all respective individual environments as the populations evolved solely in those single stressors. When present, the order of adaptation to environmental components was biased primarily towards antifungal resistance, which often involved negative fitness consequences towards other stressors (heat in particular). Although fluconazole did not impose the strongest selection of the four individual stressors, its resistance is easily acquirable due to a simple genetic basis. Interestingly, the next-most prioritized adaptation was salt resistance, which did impose the greatest selection strength, but is a highly polygenic trait (Dhar et al. 2011). This indicates that the ease of mutational accessibility to antifungal resistance drives early adaptation and trade-off dynamics, but the strength of selection determines adaptive dynamics among treatments in which the basis of adaptation are more polygenic, both consistent with mutations of greater effect size fixing first in other evolution experiments (Elena and Lenski 2003; Lang et al. 2013).

While resistance to starvation and particularly heat showed the greatest negative trade-offs at the midpoint of the experiment, these trade-offs were often nullified or even reversed by the end of the experiment. Interestingly, in the cases where we observed the greatest compensatory evolution (i.e. where maladaptation was entirely negated or even reversed), mutations that were gained or lost did not necessarily have a direct connection to the stressor in which compensation was found. Rather, affected genes were often related to a stressor to which the line was already adapted, supporting models that predict most adaptation signatures are linked to compensatory evolution rather than novel function acquisition (Pavlicev and Wagner 2012; Zhang et al. 2024).

Despite the abundance of trade-offs, some populations still managed to adapt to multiple stressors simultaneously. While a scan of first-step mutations in yeast have shown that mutations which simultaneously increase fitness across multiple traits is possible (Li, Petrov, and Sherlock 2019), this does not seem to be the primary mode of initial adaptation in our lines. Though uncommon, some lines evolved in complex environments even showed fitness surpassing that of specialist (1D) lines in at least one of the respective environments (Figure S2), indicating that some mutations had strongly positive pleiotropic effects in complex environments. Previous studies in DNA bacteriophages have shown that when selection in complex environments is particularly strong, traits constrained by antagonistic pleiotropy may acquire mutations which enhance multiple traits concurrently (McGee et al. 2016), thus overriding the trade-off costs associated with a complex selective environment. As many of these instances of adaptation greater than the specialist occur in either the 3D or 4D treatments, these populations may be forced to exploit mutations that have enhanced positive pleiotropy under the threat of extinction. While not quantified during experimental evolution, several lines at higher complexities (3D and 4D) experienced extinction at the beginning of the experiment and were restarted.

### Trade-off evolution of novel and ancestral environments

Trade-offs in non-selected (“novel”) environments were less prevalent and showed different, sometimes non-reciprocal, patterns across environments. For example, by the end of the experiment, the average fitness across all four replicate lines in each of the seven treatments containing fluconazole showed either no adaptation or maladaptation when exposed to novel fluconazole. Yet, the average fitness across replicates in five of the seven lines containing fluconazole exhibited latent adaptation to novel heat stress, including the line exposed to fluconazole alone. This asymmetry likely reflects the specificity of adaptation to each environmental component. Fluconazole adaptation in our lines commonly occurred through mutations in PDR genes, which specifically affect the efflux of azoles. However, fluconazole’s downstream effects disrupt cell wall stability (Spampinato and Leonardi 2013). Adaptation to the other stressors (salt, heat, and starvation) can involve changes to cell wall permeability and stability (Bermejo et al. 2010; Piper 1995; Hohmann 2002), and adaptive mutations for these stressors may thus have antagonistic pleiotropic effects for fluconazole resistance. Contrary to the negative pleiotropic effects found with fluconazole, the other three non-selected environments showed largely positive pleiotropy, likely because adaptations to these stressors share stress response mechanisms that provide cross-protection.

In a previous experimental evolution study, *E. coli* evolved in the presence of multiple concurrent antibiotics were more likely to acquire pleiotropic mutations that allowed them to gain novel resistances to previously unexposed antibiotics (Karve and Wagner 2022). However, contrary to these results, we observed that higher degrees of environmental complexity were associated with reduced fitness to novel stressors. This discrepancy may again result from the specificity of resistance mechanisms between the diverse stressors used in our study (salt, heat, starvation, antifungal) versus different types of antibiotics, which in contrast often involve more limited mechanisms such as efflux pumps or cell permeability changes.

Local adaptation may result in the maladaptation of a population to its ancestral environment. Indeed, trade-offs in the ancestral environment were prevalent, and also varied across levels of environmental complexity. Lines adapted to single stressors generally exhibited increased fitness in the ancestral environment and maintained these gains through the end of the experiment. In contrast, lines adapting to multiple stressors initially showed reduced fitness in the ancestral environment. While these costs often decreased by the end of the experiment, complete nullification or reversal of costs rarely occurred. This pattern aligns with empirical evidence that steeper environmental clines induce greater costs of adaptation than shallower clines (Collins and de Meaux 2009). Our observed diminishing costs of adaptation over time further support a compensatory mode of adaptation, where selection acts against antagonistic pleiotropic effects (Pavlicev and Wagner 2012).

### Conclusions

Here, we assessed both the phenotypic and genomic evolutionary dynamics in lines adapting to simple and complex environments containing multiple selective agents (full-factorial combinations of antifungal, heat, stress, and starvation). Our work highlights avenues in which patterns of adaptation and trade-off evolution can be predicted. Populations adapted rapidly to concurrent multifarious selection pressures, yet adaptive dynamics were primarily determined by the order of mutational accessibility. The acquisition of antifungal resistance typically arose first, and through remarkably similar pathways across all levels of environmental complexity. A leading strategy for the combat of both microbial infections and agricultural pests is to apply multiple antimicrobial drugs or pesticides at the same time (REX Consortium 2010). Our results indicate that unless the selection pressure from these multiple drug treatments is great enough to quickly exterminate a pathogenic or pest population, resistance will likely evolve. Further, antifungal resistance developed by similar mechanisms regardless of environmental complexity (but see (MacLean et al. 2010) for cases in which environmental context does alter the genetic basis of resistance). We also observed that evolution to complex environments led to a greater cost of adaptation to novel and ancestral environments. This suggests a concerning pattern that as populations adapt to multiple stressors simultaneously (e.g., warming temperatures and increased pollution), they may become less able to maintain high fitness under different conditions. Further, our findings indicate that as populations experience an increasing number of anthropogenic forces (Hendry, Gotanda, and Svensson 2017), their susceptibility to further habitat change may increase due to these accumulated adaptation costs. In extreme cases, habitat restoration may even decrease the fitness of a population, such as in plants adapted to heavy-metal uptake from mines (Hickey and McNeilly 1975).

Importantly, our findings indicate that adaptive dynamics in simple environments cannot be used to reliably predict outcomes in more complex ones. As environmental complexity increases, both the trajectory and genetic architecture of adaptation shift, introducing pervasive trade-offs, pleiotropic constraints, and compensatory dynamics that are rare or absent under single-stressor selection. These patterns highlight not only the need to study adaptation in ecologically realistic, multidimensional environments, but also the importance of understanding the underlying mechanisms that shape these responses. In particular, knowing whether adaptive mechanisms are stressor-specific or broadly acting will be essential for predicting the degree and direction of pleiotropy, and the constraints that emerge when multiple selective forces act simultaneously.

## Materials and Methods

### Strain information and environments

All strains used in this study were derived from a S288c background. The ancestral population was a haploid LJY170 strain (*MATɑ, hisΔ200, trpΔ63*, donated by Lars Jansen, Oxford University; see (Stajic, Perfeito, and Jansen 2019). A fluorescent strain RSY328 (*MATɑ, hisΔ200, trpΔ63, chrV 115940-116010::NatMX6-mCitrine*) was used as a reference for flow cytometry fitness measurements (see below). RSY328 was constructed by integration of a NatMX6-mCitrine cassette into an intergenic region on chromosome V, creating a deletion of the sequence spanning the positions 115,940bp to 116,010bp without affecting ORFs of neighboring genes. The NatMX6-mCitrine construct was amplified from a pRS5 plasmid, designed by insertion of mCitrine from pFA6a-ACT1-ymCitrineM233I::HISMX6 into a pAG25 backbone using HindIII and BglII restriction enzymes.

Four stressors were used in full-factorial combinations, resulting in a total of 15 environments: 0.8M NaCl, 0.5% Glucose YPD, 37°C , and 20μg/ml of the antifungal Fluconazole (Figure 1A). Populations not grown in heat stress (at 37°C) were grown at a benign 25℃. To quantify the potential selective pressure of each environment, we grew the ancestral population in each environment for 24h from an initial optical density of 0.02 at 600nm (OD600) and measured the reduction in final OD600 relative to the OD600 in the base yeast extract peptone dextrose (YPD; 1% yeast extract, 2% peptone, 2% dextrose) medium at 25°C (*n*=3) (Figure 1B).

### Experimental evolution

At the start of experimental evolution for each environment, an individual colony was isolated to serve as the ancestral population, by streaking from frozen stock culture and growing on YPD agar plates (2% agar), for 72 hours. Cells were inoculated in 5ml liquid YPD for 24 hours at 25°C and 200rpm. Subsequently, four replicates were inoculated in 5mL of each of 15 different selective environments, serving as biological replicates. Starting population size across all replicates was standardized to OD600 of 0.02 using a BioTek EPOCH2 microplate reader. Cultures were grown in their respective environments at 200 rpm. Every 48 hours 2% (100μl) of the culture was transferred to fresh media. In total, 15 transfers were completed and samples were preserved in 44% glycerol at -80℃ for phenotypic and genomic analyses (Figure 1C).

### Fitness assays

Flow cytometry competition assays were performed using a Luminex Guava easyCyte. A fluorescently labeled strain (RSY328; MATα, hisΔ200, trpΔ63, chrV 115940-116010::NatMX6-mCitrine) derived from the ancestral LJY170 background was used as a reference strain. The fluorescent strain was streaked on YPD agar plates and an individual colony was inoculated at 25°C and 200 rpm shaking for 24 h prior to measurement in the flow-cytometer. Frozen samples of the evolved populations at transfer 7 (T7) and 15 (T15) were thawed and 100μL was inoculated into 5mL of the corresponding selective environment for 24 hours. For each evolved population, three technical replicates were tested. After overnight growth, evolved populations and the fluorescent strain were mixed in a 1:1 cell ratio so that the final OD of the mixture was 0.02. Competition assays were performed in every single-stressor treatment, including the ancestral YPD medium, as well as in the relevant multi-stressor treatment. Prior to measurement with flow cytometry, mixed cultures were diluted to an OD of 0.005 and suspended in PBS (pH 7.4) solution. The count of ancestral fluorescent and non-fluorescent evolved cells were measured using flow cytometry after the initial mixing (0 hours) and after 48 hours to determine the change in proportion of evolved to ancestral cells (Supplementary Figure 9).

The relative fitness (selection coefficient) of the evolved strain to the ancestor was calculated as the slope of the linear regression of the log ratio of cell numbers 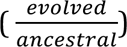 and the number of cell doublings between time points (0 hours and 48 hours). To account for any possible fitness effects of the integrated fluorescent construct, we normalized the selection coefficient values by the difference in fitness between fluorescent and non-fluorescent ancestor strains for each environment.

### Sequencing and bioinformatic processing

To culture cells for sequencing, 100μL of the frozen evolved population were inoculated in 5ml of the corresponding environment for 24 hours with shaking at 200 rpm. Approximately 1 ml of the culture was used for DNA extraction. DNA was extracted from the ancestral stock population and whole population samples at T7 and T15 of experimental evolution. The isolation of gDNA was conducted either by using a column-based method following manufacturer’s instructions (NucleoSpin Microbial DNA kit, Macherey-Nagel, Düren, Germany) or an automated system (King Fisher Duo Prime System, Thermo Scientific, MA, USA) in combination with a magnetic bead-based method (Mag-Bind Blood & Tissue DNA HDQ kit, Omega Bio-Tek, GA, USA). Quantification of gDNA samples was conducted using a fluorometric method (Qubit, Thermo Scientific, MA, USA) and library preparation was conducted either in the laboratory (DNA Prep Tagmentation kit and IDT (UD) indexes (Illumina, Hayward, CA, USA) or at the National Genomics Infrastructure Sweden (NGI) (DNA Flex library kit, Illumina, Hayward,CA, USA). The libraries were sequenced at the NGI on an Illumina MiniSeq platform with a 2×150 bp paired-end setup using a MiniSeq Mid Output kit or on an Illumina NovaSeq6000. Estimated average genome-wide coverage per sample ranged from 37.29–416.66x. Sequence data are submitted to ENA (accession number to be released upon acceptance).

The quality of sequencing reads after trimming with Trimmomatic v0.36 were assessed with FastQC v0.11.9 and MultiQC v1.11. The reference strain used was S288c (R64, GCF 000146045.2). Reads were aligned using BWA MEM v0.7.17 with the following parameters: -k 20 -r 1.3 -M. Reads were sorted and merged with samtools v1.17 (Danecek et al., 2021), and deduplicated with Picard v2.23.4 (Broad Institute). To call variants, we used GATK v4.1.4.1 HaplotypeCaller with best practices (McKenna et al., 2010). The raw VCF was filtered with the following criteria: QD < 0.5, QUAL < 20. Regions of the genome which were not uniquely mapping to itself were then filtered out using GenMap v2.4.1 (K31, E2) (Pockrandt et al., 2020), retaining 3070 variants. Sites were further filtered out if: i) variants existed in the ancestral population at <5% frequency in all populations and time-points; ii) had less than 20 reads in greater than 20% of the total samples; iii) had more than 10% of samples missing data, or iv) were found in more than 10% of samples. After filtering, 1048 variants remained. These variants were subsequently manually inspected in the Integrative Genome Viewer v2.16.0. Sites mapping to multiple locations, and homopolymeric or tandem repeat regions were discarded, resulting in a final set of 228 SNPs and 57 INDELs. Allele frequencies were calculated using the ratio between alternate and total allele depth. For each mutation, we estimated its putative effect size using snpEff (Cingolani et al., 2012), which uses the annotation of the reference strain to predict functional effects based on the type of mutation observed (e.g. synonymous, missense, frame-shift, etc).

Mutations in genic regions were associated with GO terms from the *Saccharomyces cerevisiae* Ensembl BioMart database using the R library package biomaRt (v.2.42.1). If the GO term of a mutation was found to be a lower-level GO term under one of these parent GO terms, it was grouped in the parent GO term category. Mutations which fell under multiple parent GO terms were assigned to each associated category. We then manually verified from gene entries in the Saccharomyces Genome Database (Cherry et al. 1998) that GO terms were accurately assigned to mutations for the four main GO term groups which characterize our selective treatments: osmotic stress (GO:0006970), response to starvation (GO:0042594), drug response (GO:2001023), or response to heat (GO:0009266).

### Copy number variation

We assessed potential partial- and whole-chromosome copy number variations (CNVs) in each evolved population. For partial CNVs, we calculated relative depth per population across 5kb non-overlapping windows compared to mean depth for each chromosome individually. Sites which differed by more than 20% of the mean relative depth per chromosome in the ancestral population were excluded for analyses as likely repetitive regions. Some samples had non-uniform read depth across chromosomes, where the centers of chromosomes had lower depth than the ends, leading to a wide U-shape pattern. This is a common pattern described in previous work, but is not associated with average sample coverage, flow-cell, or sequencing machine (Abbey et al. 2014; Ament-Velásquez et al. 2022; Sun et al. 2023). For every sample we fit a LOWESS curve for each chromosome based on relative depth by distance-to-chromosome-end. Relative depths were divided by the output of the LOWESS function to normalize this U-shaped pattern in the data. To detect whole-chromosome aneuploidies, we calculated relative mean depth per chromosome compared to the mean whole-genome read depth. We identified putative CNVs as windows or chromosomes which had a change in relative depth of greater than 20% from the ancestor to either subsequent time point (T7 or T15).

## Data Availability

Data is available at Figshare (doi upon release:10.6084/m9.figshare.29149616). The raw read data will be available at the European Nucleotide Archive upon acceptance. All scripts and data fall under a CC0 1.0 Universal (CC0 1.0) license.

## Code Availability

Peer review link to code and data: https://figshare.com/s/6dd76de272b899ce77f0

## Acknowledgements

We would like to acknowledge Mario Walthert and Ciaran Gilchrist for assisting in growth-curve and flow cytometry measurements. The computations and data handling were enabled by resources in project [NAISS 2023/23-487] provided by the National Academic Infrastructure for Supercomputing in Sweden (NAISS) at UPPMAX. This work was supported by the Swedish Research Council (2017-04963 and 2022-03427 to RS) and the Knut and Alice Wallenberg Foundation (2017.0163 to RS). CB is supported by “la Caixa” Foundation (ID 100010434), under agreement (LCF/BQ/AA16/11580051), by the Fonds de recherche du Québec - Nature et technologies (FRQNT) (#274987) and by PROTEO (Regroupement québécois de recherche sur la fonction, l’ingénierie et les applications des protéines) - FRQNT international internship (#293829) scholarships.

## Author contribution statement

DS and AR conceived the study and conducted the experiment. AR, DS, CB, SR, and MPCM collected the data. AR and DS analyzed the data. AR, DS, and RS wrote the paper with input from all authors.

## Supplementary Materials

**Supplementary Table 1.**
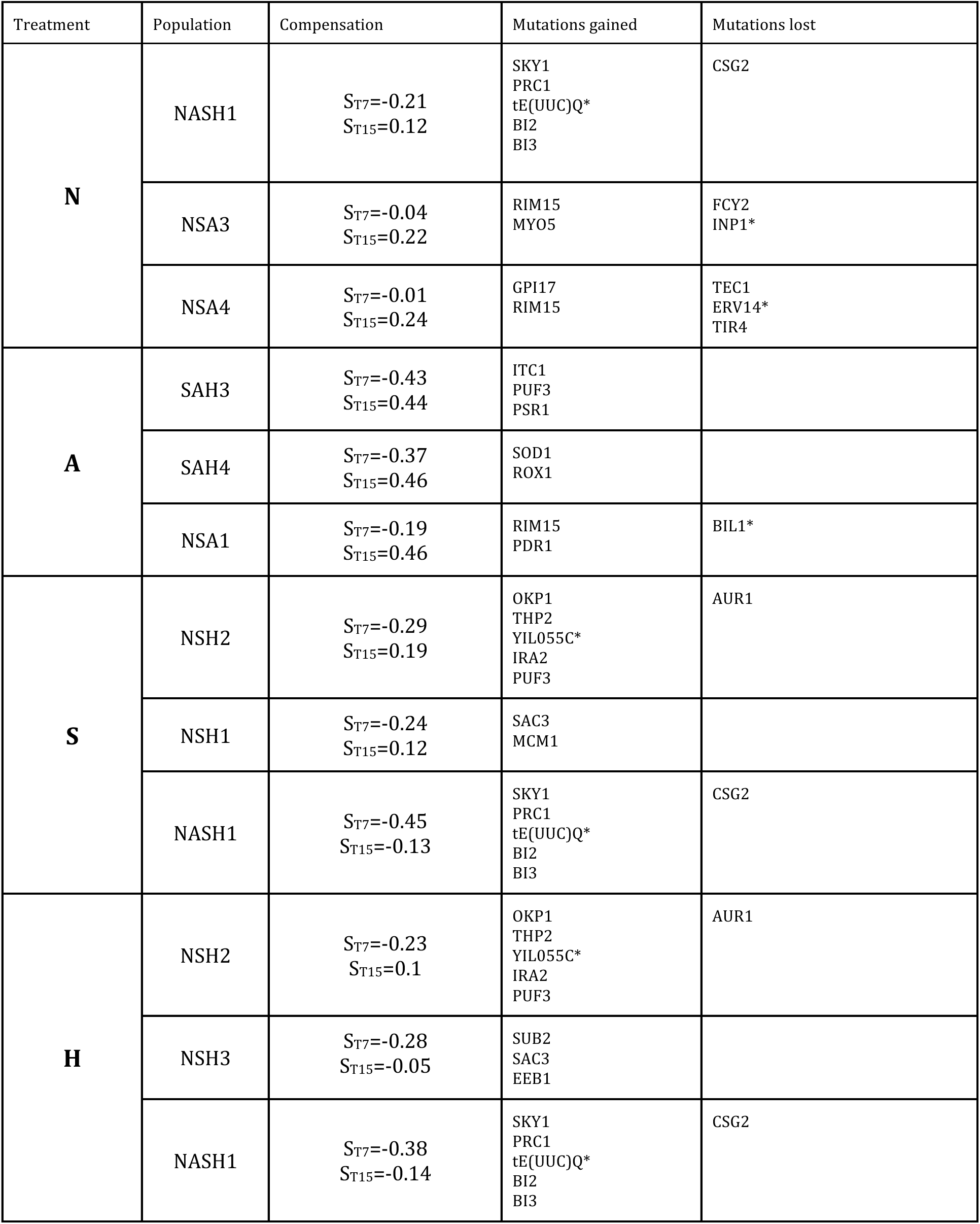
Gene mutations gained or lost during compensatory evolution (from T7 to T15). The three lines with the greatest compensatory changes (where S_T7_ <0) in fitness are shown per individual stressor. Gains and losses of gene mutations are shown in green and red, respectively. Asterisks designate upstream gene mutations not found in the protein coding region.

| Treatment | Population | Compensation | Mutations gained | Mutations lost |
| --- | --- | --- | --- | --- |
| **N** | NASH1 | $S_{T7}=-0.21$<br>$S_{T15}=0.12$ | SKY1<br>PRC1<br>tE(UUC)Q*<br>BI2<br>BI3 | CSG2 |
| | NSA3 | $S_{T7}=-0.04$<br>$S_{T15}=0.22$ | RIM15<br>MYO5 | FCY2<br>INP1* |
| | NSA4 | $S_{T7}=-0.01$<br>$S_{T15}=0.24$ | GPI17<br>RIM15 | TEC1<br>ERV14*<br>TIR4 |
| **A** | SAH3 | $S_{T7}=-0.43$<br>$S_{T15}=0.44$ | ITC1<br>PUF3<br>PSR1 | |
| | SAH4 | $S_{T7}=-0.37$<br>$S_{T15}=0.46$ | SOD1<br>ROX1 | |
| | NSA1 | $S_{T7}=-0.19$<br>$S_{T15}=0.46$ | RIM15<br>PDR1 | BIL1* |
| **S** | NSH2 | $S_{T7}=-0.29$<br>$S_{T15}=0.19$ | OKP1<br>THP2<br>YIL055C*<br>IRA2<br>PUF3 | AUR1 |
| | NSH1 | $S_{T7}=-0.24$<br>$S_{T15}=0.12$ | SAC3<br>MCM1 | |
| | NASH1 | $S_{T7}=-0.45$<br>$S_{T15}=-0.13$ | SKY1<br>PRC1<br>tE(UUC)Q*<br>BI2<br>BI3 | CSG2 |
| **H** | NSH2 | $S_{T7}=-0.23$<br>$S_{T15}=0.1$ | OKP1<br>THP2<br>YIL055C*<br>IRA2<br>PUF3 | AUR1 |
| | NSH3 | $S_{T7}=-0.28$<br>$S_{T15}=-0.05$ | SUB2<br>SAC3<br>EEB1 | |
| | NASH1 | $S_{T7}=-0.38$<br>$S_{T15}=-0.14$ | SKY1<br>PRC1<br>tE(UUC)Q*<br>BI2<br>BI3 | CSG2 |

**Supplementary Figure 1.**
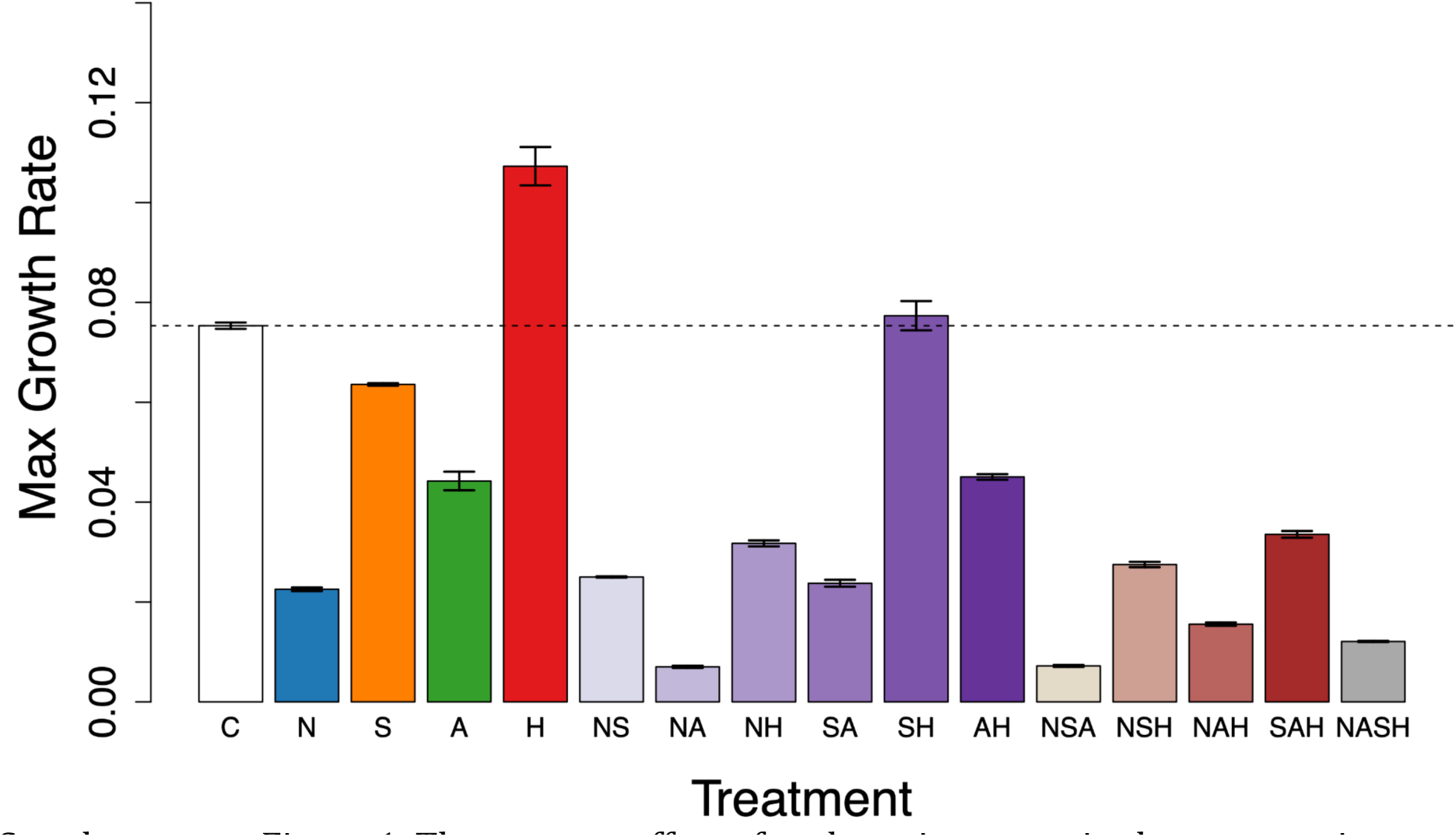
The stressor effect of each environment in the ancestor is measured as the difference in maximum growth rate in a selective environment relative to benign YPD medium. The mean estimate is given and error bars represent the standard error.

**Supplementary Figure 2.**
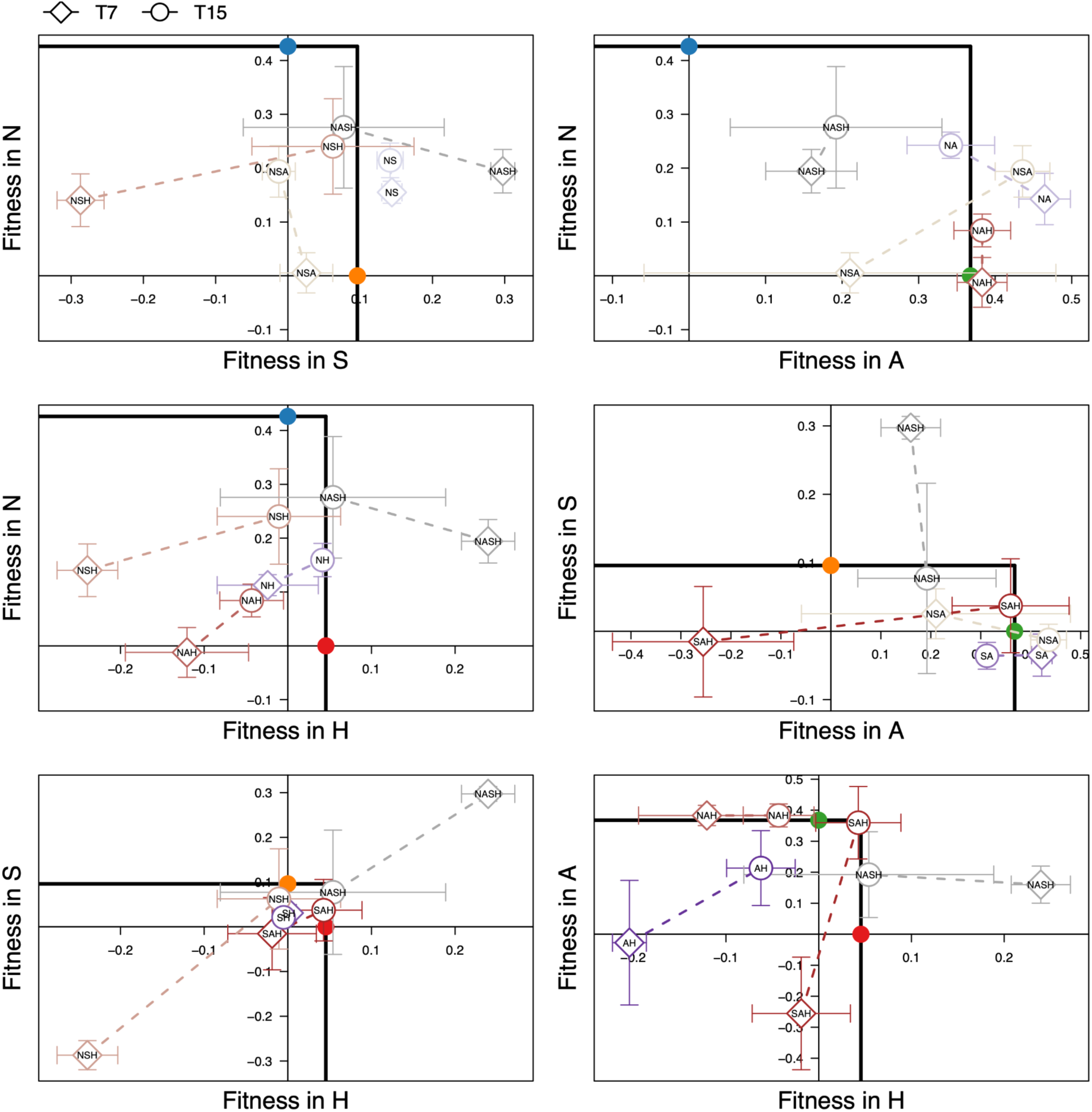
Trade-off trajectories of populations adapted to complex environments visualized for every two-way combination of individual environmental components. For each panel, the means of each line evolved in those two environmental components are shown as diamonds or circles for T7 and T15, respectively. Standard errors for selection coefficients are shown as bars for the respective axes (see Figure S1 for individual treatments). Maximum observed fitness of single environment specialists is shown as colored points projected along the respective axes. The solid black lines represent the space in which complex environments are expected to fall (i.e. less adapted in individual environments than either individual-environment specialist).

**Supplementary Figure 3.**
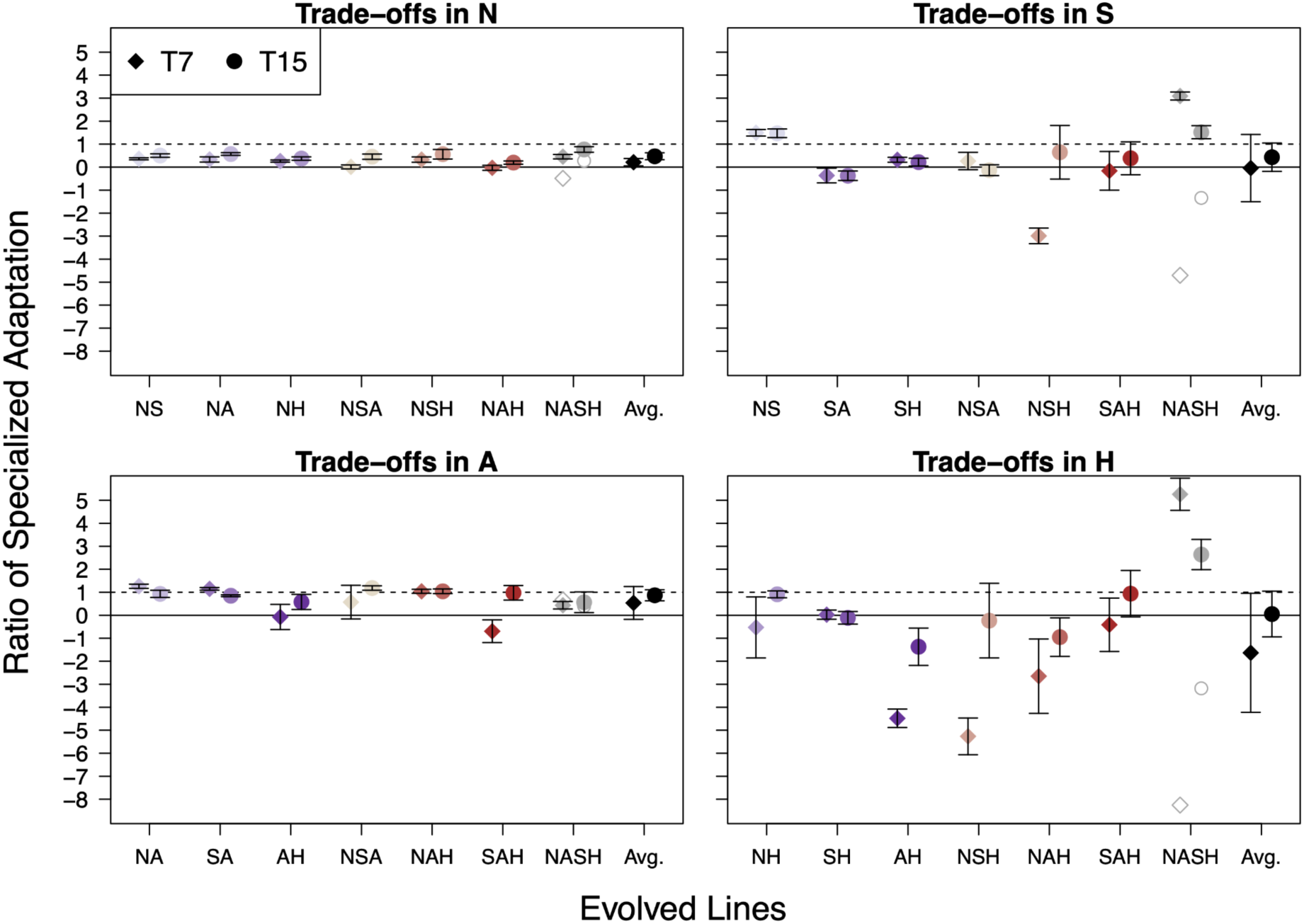
Trade-offs in each individual component of complex environment treatments. Adaptation to the individual environment by the 1D specialist are shown as projections along the respective axes. Within the lower-left of the black dotted lines represents the space in which complex environments are expected to fall (i.e. less adapted in single-environments than either 1D specialist). Means are shown as diamonds or circles for T7 and T15, respectively. Standard deviations for selection coefficients are shown as bars for the respective axes.

**Supplementary Figure 4.**
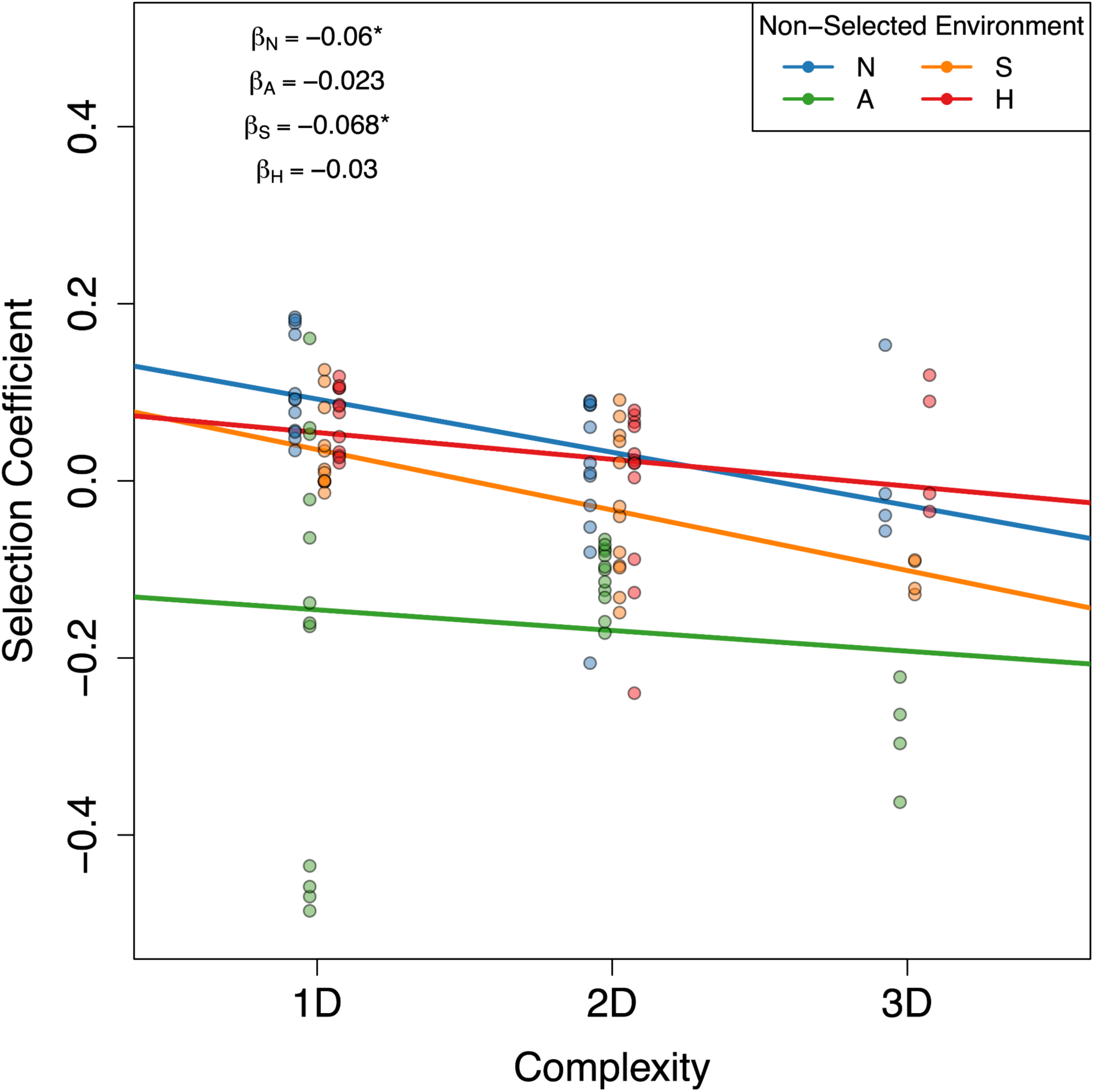
Fitness declines with environmental complexity in non-selected environments. Selection coefficients (i.e., relative fitness to the ancestor) are shown for evolved lines at T15 in test environments they were not exposed to during experimental evolution. The slope of the linear regression between fitness and complexity are shown for each non-selected environment, with significant slopes designated by an asterisk.

**Supplementary Figure 5.**
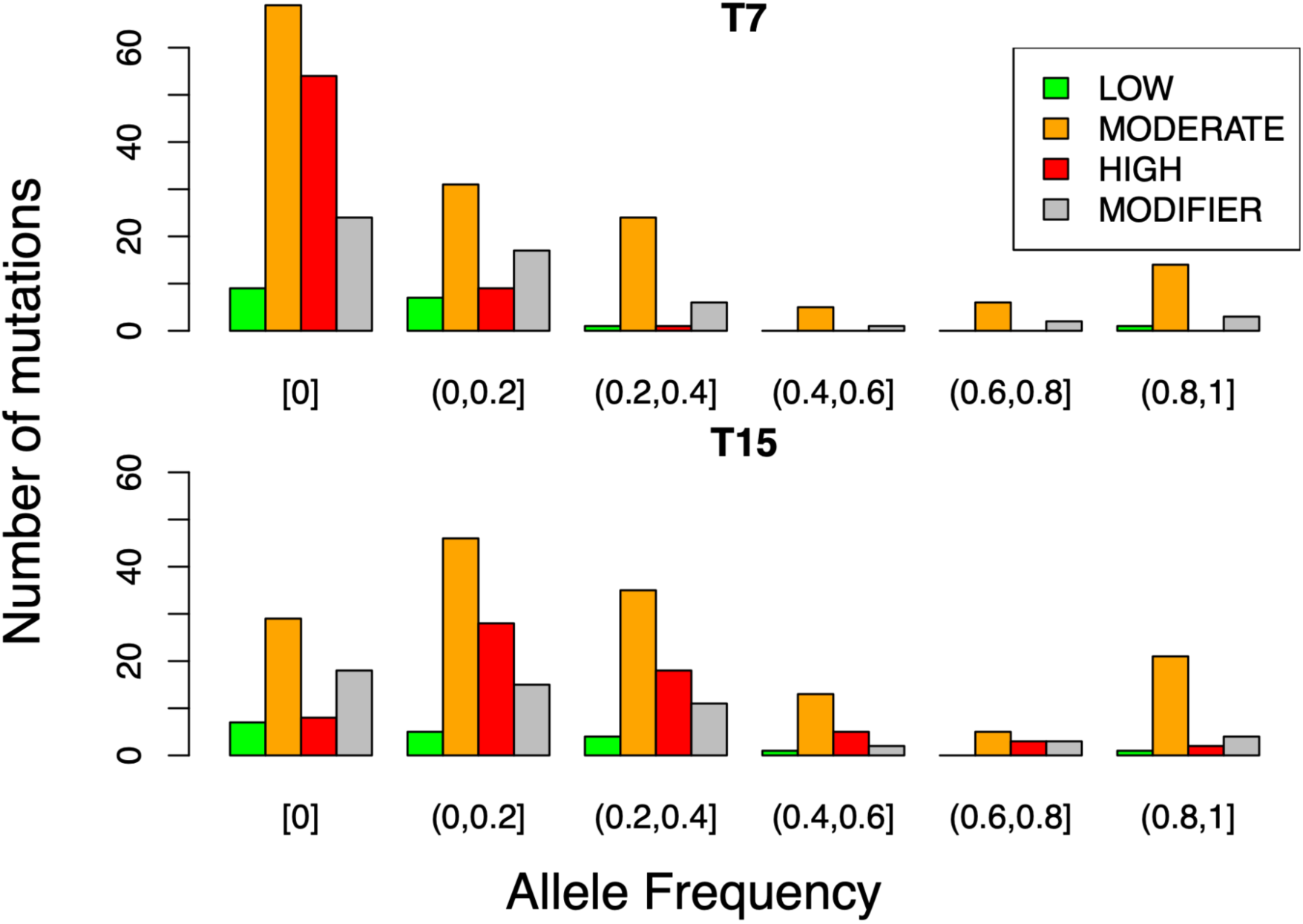
The number of types of mutations (SNPs and INDELs) across all evolved populations categorized by snpEff, binned by allele frequency. Mutations at AF=0 in T7 did not yet arise in any of the populations, and mutations at AF=0 in T15 are those that previously existed at T7 but were subsequently eliminated.

**Supplementary Figure 6.**
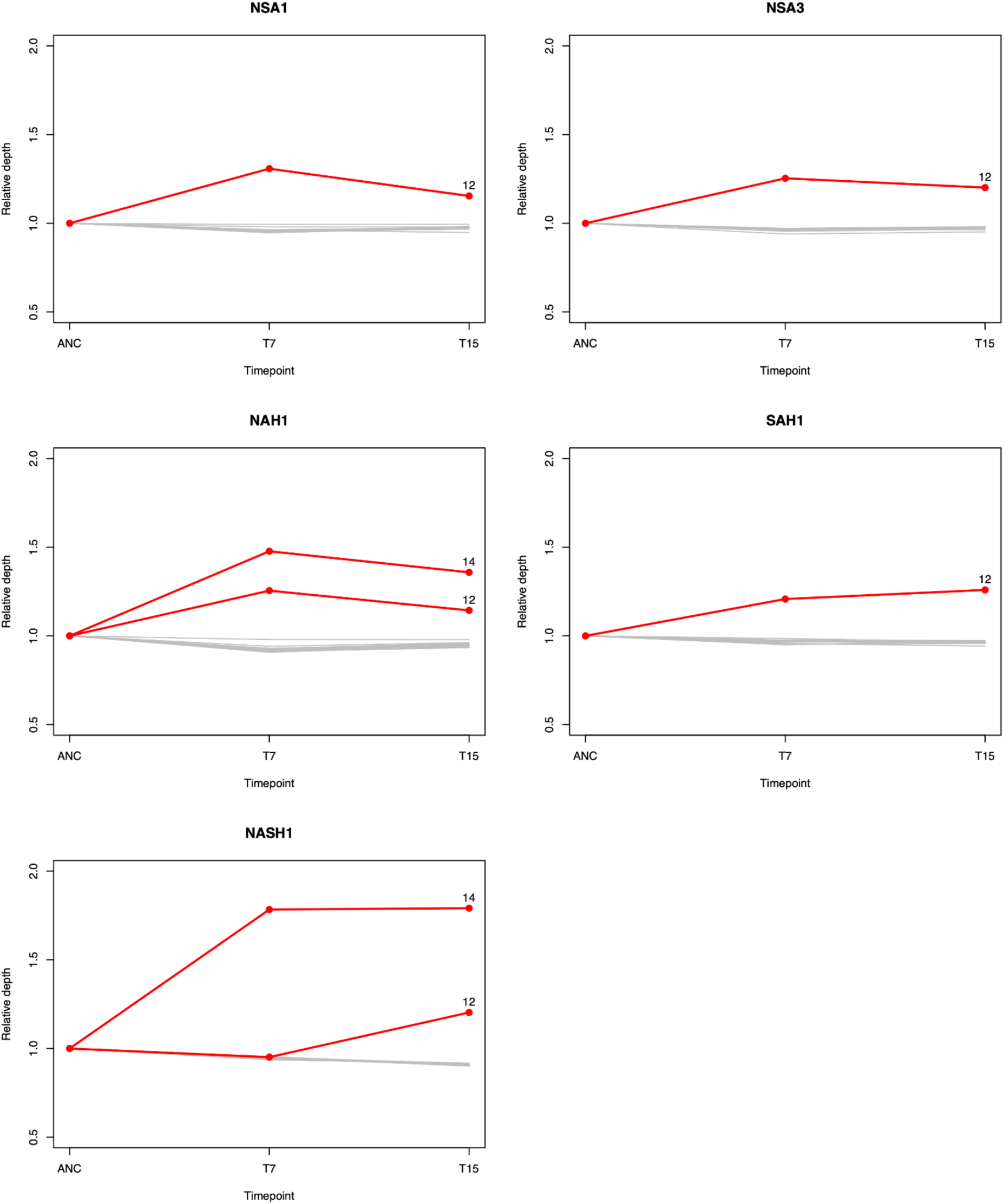
Aneuploidies identified in five populations. Plotted are the mean relative depths of each chromosome standardized to the ancestral population. In red are chromosomes which deviate from the ancestor relative depth by more than 20%. Putative aneuploidies are labeled with chromosome number.

**Supplementary Figure 7.**
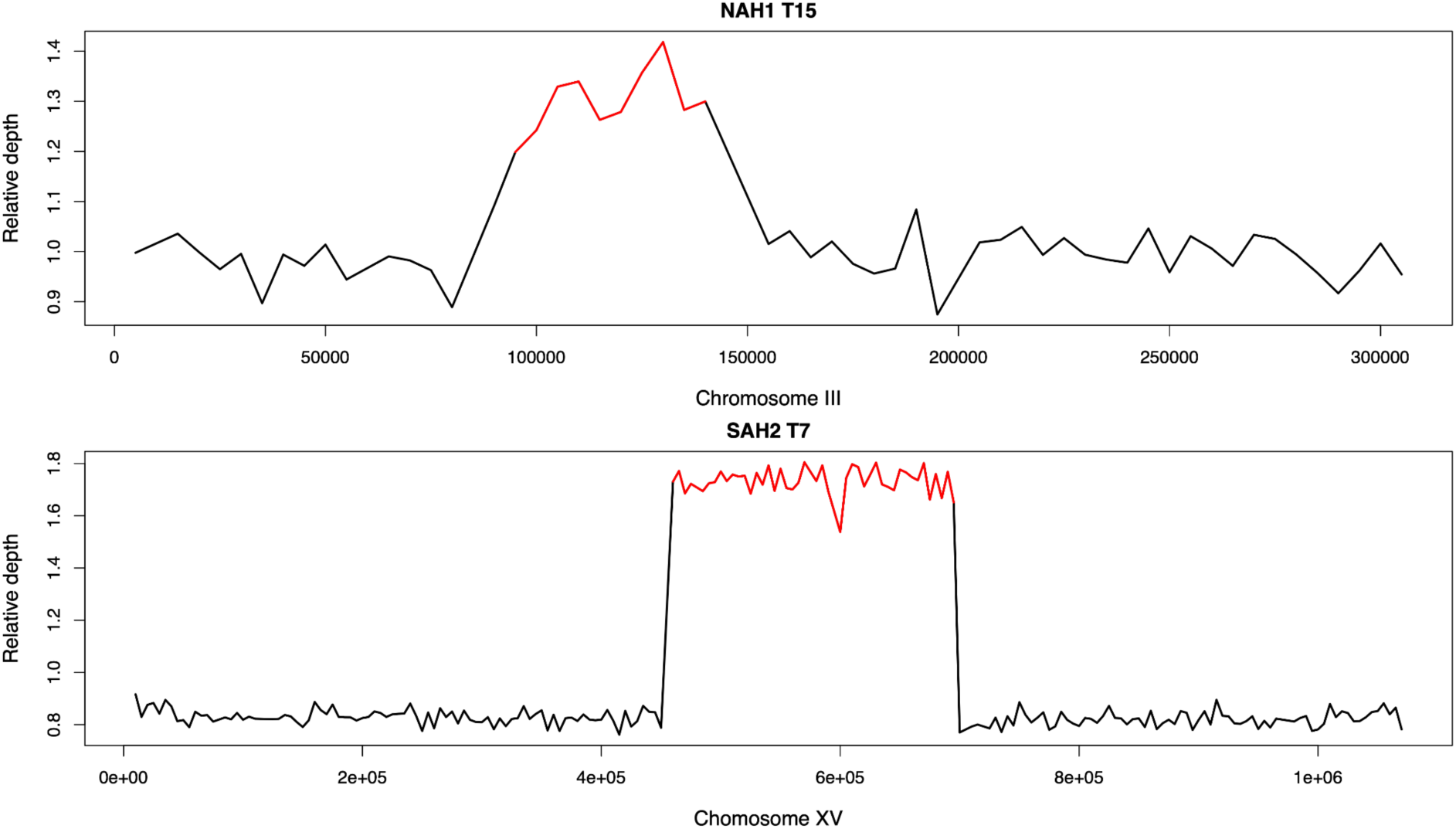
Copy number variants in two populations. Plotted are the mean relative depth across the genome in 5kb non-overlapping windows. In red are segments which deviate from the ancestor relative depth by more than 20%.

**Supplementary Figure 8.**
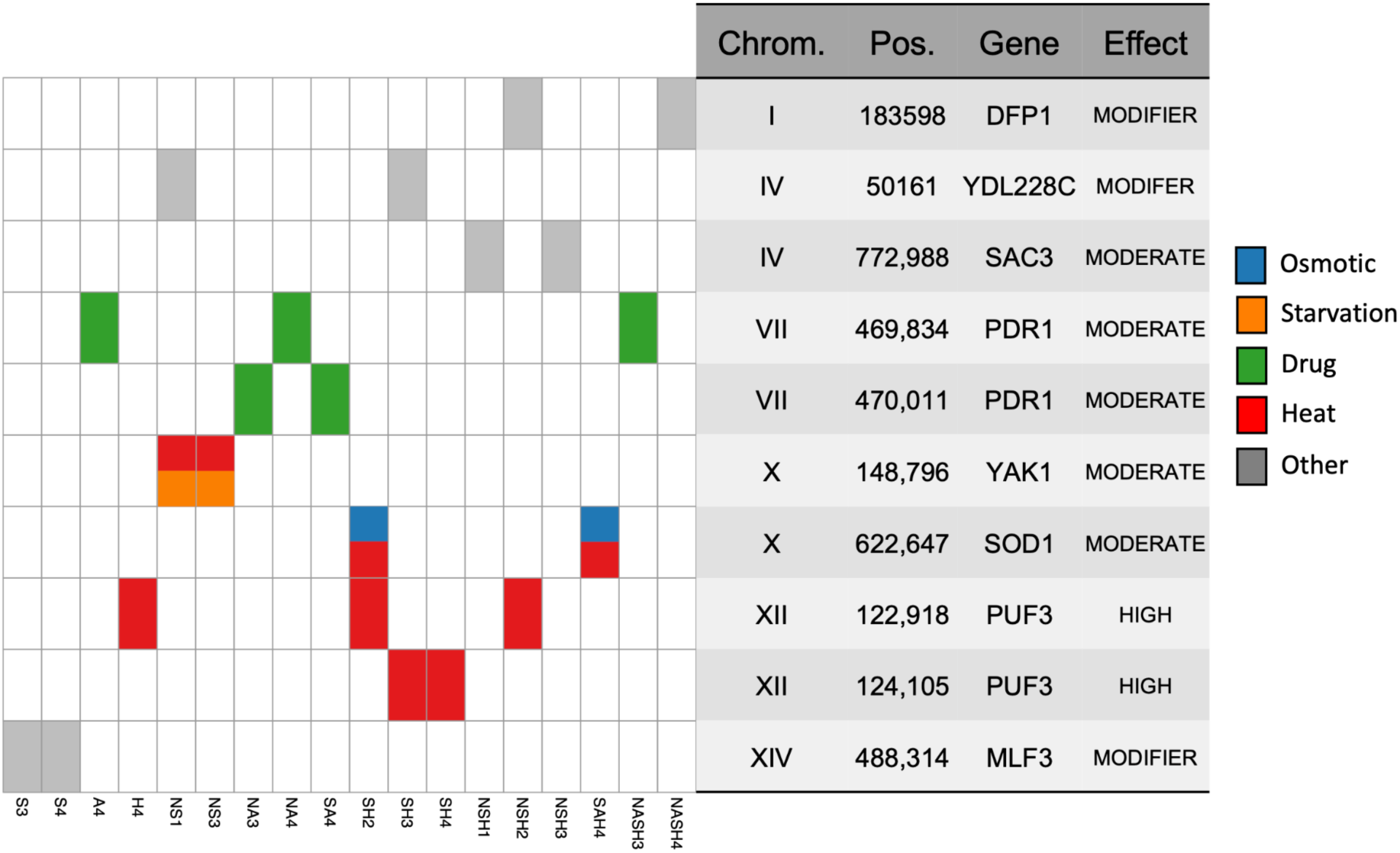
Recurrent mutations at specific SNPs across independent evolved populations. Shared mutations are colored by the parental GO category, white is no shared mutation. The table presents the chromosome, position, gene name, and predicted effect given by SnpEff for each mutation. Populations which do not contain a recurrent mutation are not shown.

**Supplementary Figure 9.**
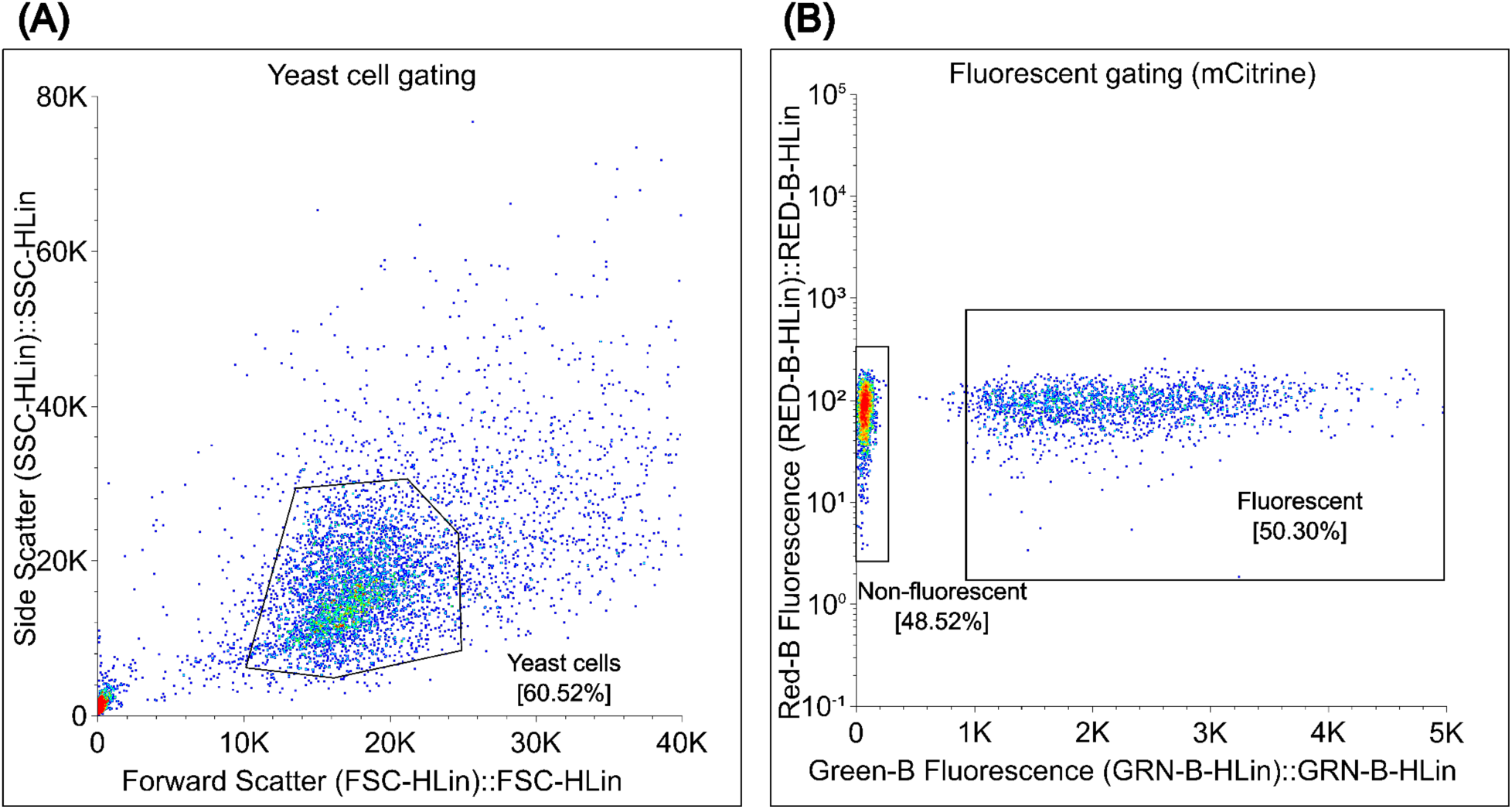
Example of gating strategy for flow cytometry. Points indicate individual events, and are colored by density (blue, low density; red, high density). A) Yeast cells were identified using a FSC gate between 10000 to 25000, and a SSC gate between 6000 to 30000. B) The proportions of fluorescent ancestral yeast to non-fluorescent ancestral or evolved lines were then discriminated using a blue 488nm laser with emission intensity (GRN-B-Hlin) greater than 1000 indicating the fluorescent ancestral cell. Note that the fluorescent ancestor and tested line appear at approximately a 1:1 ratio at 0 h of flow cytometry for this example.

